# SpecLig: Energy-Guided Hierarchical Model for Target-Specific 3D Ligand Design

**DOI:** 10.1101/2025.11.06.687093

**Authors:** Peidong Zhang, Rong Han, Xiangzhe Kong, Ting Chen, Jianzhu Ma

## Abstract

Structure-based generative models often optimize single-target affinity with ignorance of specificity, resulting in the generation of high-affinity candidates that exhibit promiscuous binding across unrelated targets. This decoupling of affinity and specificity not only compromises therapeutic efficacy but also elevates off-target risks that constrain translational potential. Therefore, we introduce SpecLig, a unified structure-based framework that jointly generates small molecules and peptides with improved target affinity and specificity. SpecLig represents a complex as a block-based graph, combining a hierarchical SE(3)-equivariant variational autoencoder with an energy-guided geometric latent-diffusion model. Chemical priors derived from block–block contact statistics are explicitly incorporated, biasing generation towards pocket-complementary fragment combinations. We benchmark SpecLig on peptide and small-molecule tasks using standard public datasets and propose precision/breadth testing paradigms to quantify specificity. Across multiple evaluations, ligand candidates generated by SpecLig usually bind to the target pocket with high specificity and affinity while maintaining competitive advantages in other attributes. Ablations indicate that both hierarchical representation and energy guidance contribute to success. Finally, we present multiple real applications that demonstrate how SpecLig improves ligands in natural complexes to mitigate potential off-target risks. SpecLig, therefore, provides a practical route to prioritize higher-specificity designs for downstream experimental validation. The codes are available at: https://github.com/CQ-zhang-2016/SpecLig.

## 1 Introduction

Designing ligands with target specificity advances both mechanistic biology and drug discovery^1, 2^. Recent deep-learning generative models^3–8^ have shifted ligand discovery from large-scale screening^9–12^ toward rational, structure-guided design. Structure-based drug design (SBDD) is central to this trend because it leverages the receptor’s three-dimensional (3D) geometry to constrain generation in chemically and spatially relevant regions of ligand space^13–15^.

Ligands have diverse chemical modalities, including small molecules, peptides, and antibodies. Each class has distinct pharmacokinetic and functional profiles (e.g., oral bioavailability for small molecules^16^; recognition motifs for peptides^17^). Despite differences in scale and physicochemical properties, ligand–receptor binding follows the same physicochemical rules, such as bond types, bond angles, lengths, and steric clashes^18, 19^. Common generative paradigms such as autoregressive models^20–22^, diffusion-based methods^23–26^, flow-matching^27–29^, retrieval-augmented generation^30, 31^, energy optimization^32–35^, iterative refinement^36–39^, and voxel-based models^40, 41^ have been successfully applied across these ligand classes. Recent works^42–46^ have considered unified generative frameworks that accommodate representational differences among ligand types.

Off-target is a well known challenge in traditional drug discovery^47–50^. Most SBDD models learn ligand distributions conditioned on a single target structure and, therefore, emphasize recurring motifs from the training data. While such motifs can increase predicted affinity, their systematic reuse can reduce target specificity. For examples (Figure 1a and b), voxbind and PepGLAD produced candidates with higher predicted affinity than native ligands for receptors (UniProt IDs: P14779 and P33247); however, they also showed enhanced binding to unrelated proteins. This decoupling of affinity and specificity indicates a systematic bias toward promiscuous binders, increasing off-target risk and constraining translational potential^51–53^. In the interaction analyses shown in Figure 1a and 1b, some fragments in the designed ligands contributed little to binding with the intended target yet dominated interactions with off-targets. We further analyzed several advanced SBDD models and categorized their outputs by specificity. As shown in Figure 1c, small molecules with low specificity tend to contain a markedly higher fraction (≈5–10%) of polar groups, which promote promiscuous binding across multiple targets. Figure 1d shows that highly specific peptides typically contain a higher fraction (≈3–10%) of helical structure, whereas non-helical flexible segments increase exposure risk. Experimental details are provided in Supplementary Chapter 3. These pervasive generative biases in deep learning models highlight the need to explicitly consider specificity in ligand design.

**Figure 1.**
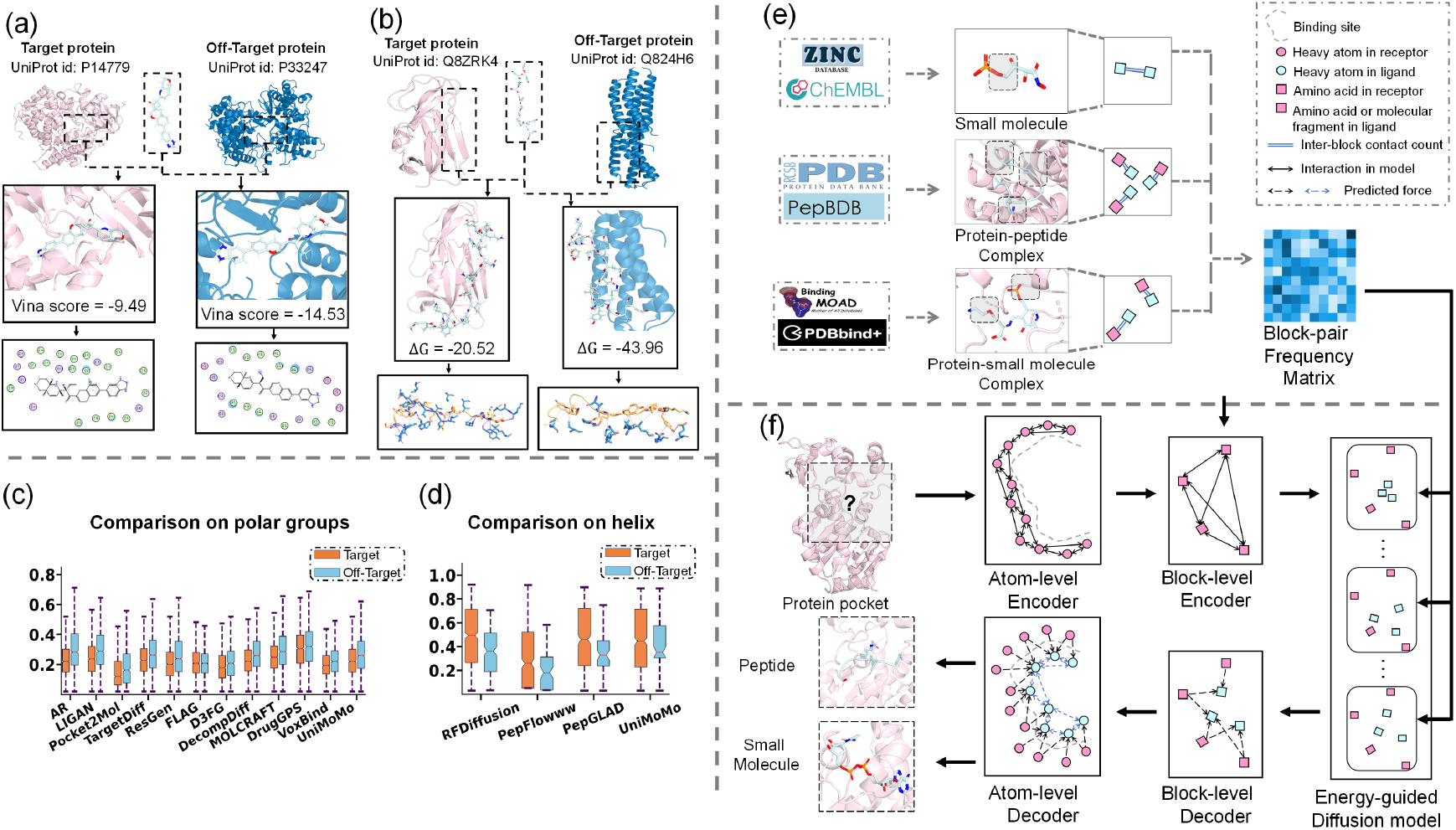
Off-target analyses and the main architecture of SpecLig. We present two representative cases in (a) and (b). Existing SBDD models generate small-molecule and peptide ligands with high predicted affinity for their intended receptors (pink) but also exhibit undesired binding to unrelated proteins (blue). (c) and (d) provide a comparative analysis of chemical and structural features distinguishing high- and low-specificity designs. (c) differences in polar-group ratios among small molecules, and (d) differences in helical content among peptides. (e) and (f) show the overview of the proposed SpecLig framework. (e) depicts the construction of the block–block frequency matrix used as a statistical energy prior, and (f) illustrates how this prior guides the latent diffusion–based generation process within the hierarchical VAE architecture of SpecLig.

Standard SBDD benchmarks focus on single-target docking scores, measuring spatial and electrostatic complementarity^54, 55^. Generative models can produce candidates with high docking scores through excessive modifications, masking potential promiscuity. Prior attempts^56^ to quantify specificity in small-molecule design by comparing against a random non-target are limited by randomness and provide little actionable guidance. To address this, we design a suite of specificity metrics that quantify binding preferences across multi-target contexts and are applicable to both small-molecule and peptide design. Using these metrics, we find that many model-generated ligands with improved single-target affinity exhibit reduced specificity relative to reference ligands. It is consistent with the empirical difficulty of improving selectivity versus affinity^57^.

We think that achieving target specificity requires moving beyond single-structure conditioning to incorporate evolutionary binding preferences. We address this by introducing energy-guided diffusion that leverages statistical contact frequencies between molecular fragments derived from native protein-ligand complexes. Unlike conventional energy functions based on physical potentials, our potential terms quantify the empirical preference for specific inter-fragment interactions across diverse targets. Based on this, we propose SpecLig (shown in Figure 1), a unified generative framework that integrates structure-based design with statistical energy guidance to simultaneously design high-specificity small molecules and peptides. SpecLig employs a hierarchical graph neural network (GNN): an atom-level encoder for local chemistry and bond order, and a block-level encoder that represents residues or predefined molecular fragments to capture global topology with reduced cost. Fragments are derived by subgraph partitioning^58^ of a molecular library to form a reusable vocabulary. Crucially, chemical priors derived from native complexes are incorporated both as features in training and as additive guidance during latent diffusion sampling. It biases generation toward a chemical space that is more likely to enhance specific binding under the given conditions, improving binding affinity while preserving geometric accuracy.

We evaluated SpecLig on both peptide and small molecule benchmarks. The results show that our unified framework attains competitive performance on standard benchmarks while consistently producing ligands with higher specificity. Moreover, we demonstrate that SpecLig can simultaneously optimize affinity and specificity for native ligands that exhibit potential off-target risks, illustrating its practical utility in advancing candidate molecules toward safer leads.

## 2 Methods

SpecLig addresses the critical challenge of binding specificity in structure-based ligand design through statistical energy guidance. SpecLig models protein–ligand complexes as graphs of molecular fragments (blocks) and combines a hierarchical variational autoencoder (VAE^59^) with an E(3)-equivariant latent diffusion model. A hierarchical encoder extracts multi-scale geometric representations, and a decoder with a similar architecture reconstructs block types and full-atom 3D coordinates. The core innovation lies in an energy-guided diffusion process that leverages empirical block–block interaction statistics from natural complexes, steering generation toward target-specific binding configurations while maintaining high affinity. Supplementary Chapter 2 discusses related work on SpecLig.

### 2.1 Notations and overview of SpecLig

We denote a protein–ligand complex as a block-based graph *G* = (*V, E*). Each node, *v*_*i*_ ∈ *V*, is an unordered set of atoms represented as 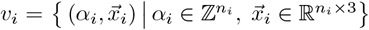, consisting of all the element types and 3D coordinates for the *n*_*i*_ atoms in the *i*th block. Edges *E* record intra- and inter-block chemical bonds and spatial adjacency. A block vocabulary *S* (canonical amino-acid residues and predefined small-molecule fragments) assigns each block a type *s*_*i*_ ∈ *S*. For controllable generation, we introduce a flag *p*_*i*_ ∈ {0, 1} in each block: *p*_*i*_ = 1 restricts sampling to canonical residues; otherwise, it samples from the full vocabulary. The pre-computed empirical block–block frequency matrix is denoted as 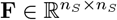, where *n*_*s*_ is the length of *S*.

For conditional design, the pocket and the ligand are represented as *G*_*P*_ and *G*_*L*_, and the task is to model the conditional generation distribution *p*(*G*_*L*_ *G*_*P*_). Assuming that *ξ, ϕ*, and *θ* respectively represent the parameter sets of the VAE encoder, decoder, and latent diffusion model, the generation probability can be expressed in marginalized form:

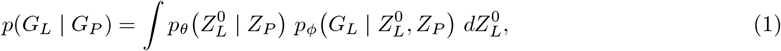

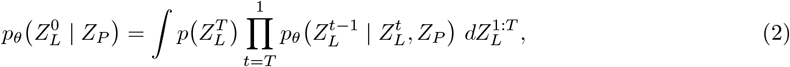

In Equations 1 and 2, *Z*_*P*_ denotes the latent representation of the pocket, 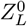 denotes the noise-free latent representation of the ligand, and 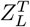 is the terminal latent state approaching the prior (Gaussian noise). Finally, the pocket is defined as blocks whose reference points (*C*_*β*_ or the fragment centroid) lie within 10Å of any native ligand atom, preventing information leakage^21^.

### 2.2 Implementation of hierarchical VAE

We decompose the VAE into hierarchical modules ℰ_*ξ*_ = {ℰ_*ξ*,1_, ℰ_*ξ*,2_} and 𝒟_*ϕ*_ = {𝒟_*ϕ*,1_, 𝒟_*ϕ*,2_}. All networks are implemented with SE(3)-equivariant transformers^60^ to guarantee translation–rotation equivariance.

#### Atom-scale to block-scale encoding

The atom-scale encoder *E*_*ξ*,1_ encodes each atom with its element type, parent block type, canonical-residue flag, and chain id. We augment atom features with a correlated projection learned from the empirical frequency matrix **F** after temperature-scaled normalization.

As shown in Figure 1f, the information flow is restricted to operate within *G*_*L*_ and *G*_*P*_ separately when constructing the k-nearest neighbors (KNN) graph. Edge features are defined as {*e*_*ab*_, **d**_*ab*_, ***β***_*ab*_ }, indicating whether the two atoms belong to the same block, their relative distance, and any candidate chemical-bond types. The block-scale encoder *E*_*ξ*,2_ constructs a coarser KNN graph on top of *E*_*ξ*,1_’s outputs. The edge features are updated to the relative distances between block centroids. Define 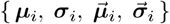 as the distributional parameters learned by the encoders. After reparameterized sampling^61^, we could obtain the block-level latent representations for *G*_*L*_ and *G*_*P*_, expressed as:

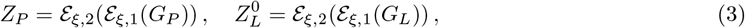

The latent representation of *i*th block could be defined 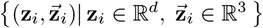, where *d* is set to 8. Before decoding, Gaussian noise is injected to model the deviations encountered during the latent diffusion process, thereby improving robustness.

#### Block-scale to atom-scale decoding

The hierarchical decoders decode block types and reconstruct atomic coordinates at their respective scales. Information flow is constrained to be unidirectional from *G*_*P*_ to *G*_*L*_ (shown in Figure 1f). The block-scale decoder predicts block type probabilities and coarse block centroid positions. Outputs are normalized with softmax. Querying the block vocabulary *S* to retrieve the set of atom types, intra-block bonds, and the initial atomic positions. Specifically, design task is identified based on user input, and the block type prediction is explicitly restricted to the relevant vocabulary.

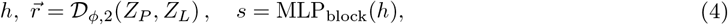

We then adopt an iterative, flow-matching–like decoding scheme^62^. For each block *i* and each time step *t*, we predict two atom-level features 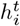 and 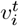, which indicate per-atom feature representations and relative coordinate updates, respectively. Initial atom positions 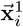 are sampled randomly around the predicted centroids. Time step *t* decreases from 1.0 to 0.0 with Δ*t* = 0.1. The update dynamics follow:

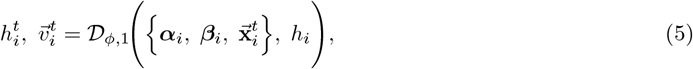

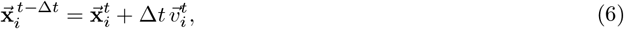

#### Chemical bond prediction

From the final atomic representations 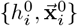 produced by the hierarchical decoder, we predict inter-block bond distributions using a two-stage procedure (shown in Equations 7 and 8). A small MLP-based frontier predictor first selects spatially-proximal (distance < 3.5Å), inter-block atom pairs. Then another MLP-based bond predictor is used.

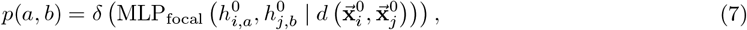

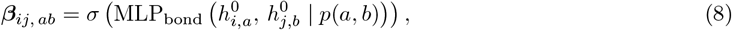

Where (*a, b*) are the indices of two different atoms in the *i*th block and the *j*th block, separately. *d*(·), δ(·), and *σ*(·) respectively indicate the distance calculation, gated function, and softmax activation. In practice, predicted bond assignments are accepted based on confidence ranking and valence compatibility. To refine geometric and chemical accuracy, the entire encoder–decoder pipeline may be iteratively run for multiple design cycles, depending on the task.

### 2.3 Energy-guided geometric diffusion model

We perform conditional diffusion in the latent space and inject a data-driven energy guidance during sampling that explicitly favors fragment pairings empirically associated with cognate binding. A vanilla SpecLig variant without energy guidance is reported in Supplementary Figure S1 as an ablation.

#### Statistical prior construction

As illustrated in Figure 1e, we precomputed empirical block–block contact priors from multiple curated sources. (i) Fragment-fragment occurrences from over a million small molecules sampled from ZINC15^63^ and ChEMBL^64^ databases, decomposed by the principal-subgraph algorithm^58^. (ii) Residue pairs bound by inter-chain hydrogen bonding from RCSB PDB^65^ and PepBDB^66^ databases, using trajectory-based hydrogen-bond analysis software. (iii) protein–ligand interaction frequencies from PDBbind^67^ and Binding-MOAD^68^ analyzed with BINANA. Source-wise frequency matrices are normalized to reduce modal bias and log-transformed to yield continuous statistical potentials. Noted that the data used to construct the frequency matrix were strictly filtered to entries published prior to 2020, ensuring no overlap with the evaluation benchmarks. Rather than memorizing specific complex structures, it serves as a coarse-grained statistical prior, conceptually analogous to BLOSUM^69^ matrices in protein sequence analysis. The frequency matrix captures universal, target-independent bonding probabilities between amino acids and molecular fragments, leveraging recurring physicochemical patterns to guide generation.

#### Why does it improve specificity

Frequent fragments in training data often reflect general potential rather than pocket specificity. Using **F** as a statistical potential reweights sampling toward fragment combinations that historically co-occur in native complexes for pockets similar to the query, thereby reducing the generation of promiscuity-prone motifs.

#### Standard diffusion process

We adopt a standard diffusion forward–backward scheme. Let the encoder outputs for the pocket and ligand (*Z*_*P*_ and *Z*_*L*_) be the clean latent variables 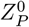 and 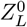, respectively. 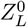 is normalized following empirical practices^70^. We apply a discrete-time Gaussian forward noising process with a cosine schedule 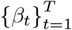, then define *α*_*t*_ = 1 − *β*_*t*_ and 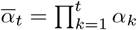. Using the attribute latents 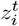 as an example, the forward Markov chain is given by:

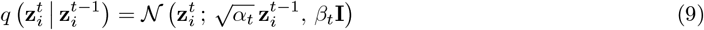

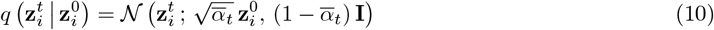

The reverse denoising is parameterized conditionally as:

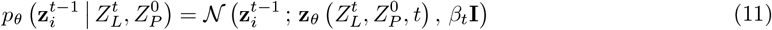

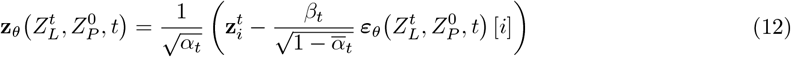

Where ***ε***_*θ*_ also uses the SE(3)-equivariant network. The latent representation of block position 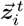 follows the same forward–backward scheme.

#### Energy integration into sampling

During sampling, we guide each reverse step with an energy term derived from the block–block bonding prior to favoring chemically plausible ligands. Given a denoised estimate 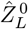 reconstructed from the current predicted noise, we decode block-type probability vectors ŝ with the frozen 𝒟_*ϕ*,2_ following Equation 4. For a block pair (*i, j*), the pairwise energy term is defined as:

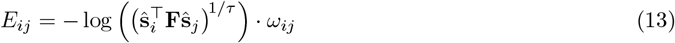

Where *τ* is a temperature smoothing factor, and *ω*_*ij*_ is a distance-dependent decay. For stability, we apply thresholding to the predicted probabilities. The total energy *E* is normalized based on the ligand’s molecular mass. Crucially, this energy term quantifies how well the current fragment combination matches the target-selective binding patterns observed in nature.

We integrate the energy into the reverse dynamics by backpropagating the energy gradient from the block probabilities into the noise space. From the reverse format of Equation 10, we could obtain the denoised representation 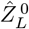 reconstructed by the current predicted noise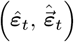. Thus, the backpropagation follows:

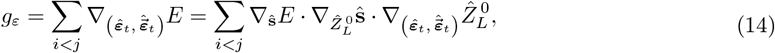

Gradient update only the noise-prediction network. Using the attribute latents as an example, we adjust the sampling process by stepping in the negative energy gradient direction:

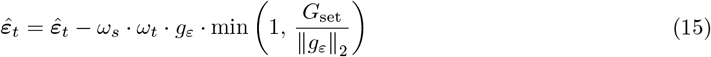

To guarantee stability, gradients are clipped by their norm to a preset bound *G*_set_, and the guidance weight *ω*_*t*_ decays with time steps. The weight *ω*_*s*_ is explicitly assigned based on the task type to modulate the overall response intensity. Details of the sensitivity analysis for *ω*_*s*_ can be found in Supplementary Chapter 13. Besides, pseudocodes for the full training and guided sampling algorithms are provided in Supplementary Chapters 4 and 5.

### 2.4 Training procedure and loss design

We design a composite loss to optimize multiple objectives. The hierarchical VAE and the latent diffusion model are trained sequentially, following best practices^42, 43^. In the hierarchical VAE stage, atom-scale loss terms include: (i) a focal atom-pair classification loss and bond-type cross-entropy for inter-block bond prediction, (ii) a mean-squared error (MSE) loss on the predicted velocity fields produced during iterative decoding, and (iii) a paired-distance loss between atom pairs. The velocity-field supervision is obtained by interpolating between noisy and ground-truth coordinates. The paired-distance loss is applied only for adjacent (≤ 6Å) atoms in the native structure and during early decoding steps (*t* ≤ 0.25).

At the block scale, we impose Kullback–Leibler (KL) regularization on both attribute latents and coordinate latents, and supervise block-type classification and coarse centroid regression. Lastly, we consider the contrastive loss to align the ligand and target pockets from a global perspective. Global descriptors are obtained by averaging per-node features. The triplet-based contrastive loss pulls the ligand descriptors closer to their corresponding pocket and pushes them away from a randomly sampled pocket. The training objective is the weighted sum of the atom-scale, block-scale, and global contrastive terms. The training uses teacher forcing: atomic types, intra-block bonds, and 50% of inter-block bonds are exposed to the VAE. Additionally, 5% of pocket residues are also masked during training.

For the latent diffusion stage, we train with a weighted sum of the denoising MSE loss and a latent perceptual loss (LPL). The LPL term follows the original formulation^71^, and the detailed implementation is provided in Supplementary Chapter 6. The overall diffusion-stage loss follows below. Lastly, we provide the default hyperparameters in Supplementary Chapter 9:

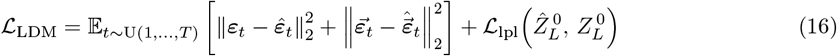

## 3 Experiments

We trained and evaluated SpecLig on peptide (PepBench^42^, ProtFrag^42^, and LNR^72^) and small-molecule (CrossDocked2020^73^) design benchmarks using standardized protocols, following the splitting recommendations provided by the authors^20, 42^. Considering the splitting bias suggested by work^74^, we provided the similarity between the training and test sets in Supplementary Chapter 1. Domain-specific baselines were trained only on their respective domain data, whereas the unified frameworks were trained on the multi-domain training data. To enhance small-molecule chemistry modeling, we augmented training with 100,000 ChEMBL^64^ compounds under 20% structural masking. All dataset details are provided in Supplementary Chapter 1.

Evaluation employed commonly used domain metrics grouped by evaluation objective. We report (i) per-category weighted ranking scores in overall comparisons and (ii) per-metric numerical results within each category. Ranking scores were computed as 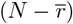, where *N* is the number of competing models in each category and *r* is the mean rank across the category’s individual metrics. Notably, the weights across metric categories in our overall comparisons were manually assigned to prioritize target specificity within the broader context of this multi-objective optimization task. All numerical results reported represent per-sample means calculated across the respective test sets. They were normalized to [0, 1] for clarity, the higher, the better.

We introduce two paradigms to quantify specificity. The precision paradigm docks designs against the target and one randomly chosen non-target using a high-accuracy engine; the breadth paradigm screens designs against a protein set that includes the target to mimic multi-target pharmacological environments. For peptides, we use PyRosetta side-chain docking (multiple precision modes, set size 100); for small molecules, we use Uni-Dock^75^ (breadth scans, set size 2000) and AutoDock Vina^76^ (precision). Finally, comprehensive ablations appear in Supplementary Chapter 10.

### 3.1 Small molecules

#### Setup

We follow prior benchmarks^77^ and group metrics into five categories: **Substructure, Geometry, Chemistry, Specificity**, and **Interaction. Substructure** measures agreement at the atom-type, ring, and functional-group levels using the Jensen–Shannon divergence (JSD, ↓) for distributional shift and per-molecule mean absolute error (MAE, ↓) for frequency deviations. **Geometry** assesses local geometric validity via the JSD of bond lengths (JSD_BL_, ↓) and bond angles (JSD_BA_, ↓), as well as atom-collision statistics: mean collided-atom fraction (Ratio_cca_, ↓) and the fraction of collided molecules (Ratio_cm_, ↓). **Chemistry** quantifies drug-likeness (QED, ↑), normalized synthetic accessibility (SA, ↓), LogP (− 0.4 ≲ ideal ≲ 5.6), and the ratio satisfying Lipinski rules (LPSK, ↑). In **Specificity**, Δ**E**_pair_ (↓) and Ratio_pair_ (↑) report the score difference between target and non-target dockings and the fraction of cases in which the target scores better, respectively; Δ**E**_mean_ (↓) and Ratio_20_ (↑) report the extent to which the target score exceeds the set mean and the ratio of designs in the top 20%. **Interaction** evaluates binding performance: mean AutoDock Vina score (E, ↓) and fraction exceeding the reference ligand (IMP, ↑). To correct for the effects of molecular size, we report the per-pocket mean relative energy gain (MPBG, ↑) and ligand binding efficiency normalized per atom (LBE, ↑). Interaction patterns are further characterized via seven PLIP-detected^78^ interaction types: population-level agreement is quantified by JSD_OA_ (↓) and MAE_OA_ (↓), per-pocket agreement by JSD_PP_ (↓) and MAE_PP_ (↓). Besides, JSD_bs_ (↓) compares the fragment composition in the binding site.

#### Baselines

We compare SpecLig to representative methods with various generative paradigms: 3D autoregressive (AR^20^, Pocket2Mol^21^, ResGen^22^), diffusion-based (TargetDiff^24^, DecompDiff^25^), fragment-based (FLAG^36^, D3FG^37^, DrugGPS^39^), voxel-based (LiGAN^40^, VoxBind^41^), and continuous-space approaches (MolCRAFT^79^, UniMoMo^43^). Baseline designs follow the CBGBench and MOLCRAFT benchmarks where available; others use standard reported settings.

#### Results and discussion

Table 1 and Figure 2 summarize weighted scores and per-metric ranks (complete results in Supplementary Chapter 7). SpecLig exhibits robust gains across multiple categories. On Specificity, it ranks first or second across metrics (Δ**E**_pair_ = − 0.83, Ratio_pair_ = 58.73%, Δ**E**_mean_ = − 0.75, Ratio_20_ = 30.17%, Supplementary Table S3), indicating preferential binding to intended pockets. In Interaction metrics, SpecLig achieves near-optimal mean Vina scores and the highest MPBG (= 15.17), representing a 53.4% relative gain versus VoxBind (Supplementary Table S5). Chemistry and Substructure metrics remain competitive, implying improved pocket specificity without loss of drug-likeness or synthetic accessibility. Geometrically, collision rates are low, though bond-length distributions still show room for refinement.

**Table 1.**
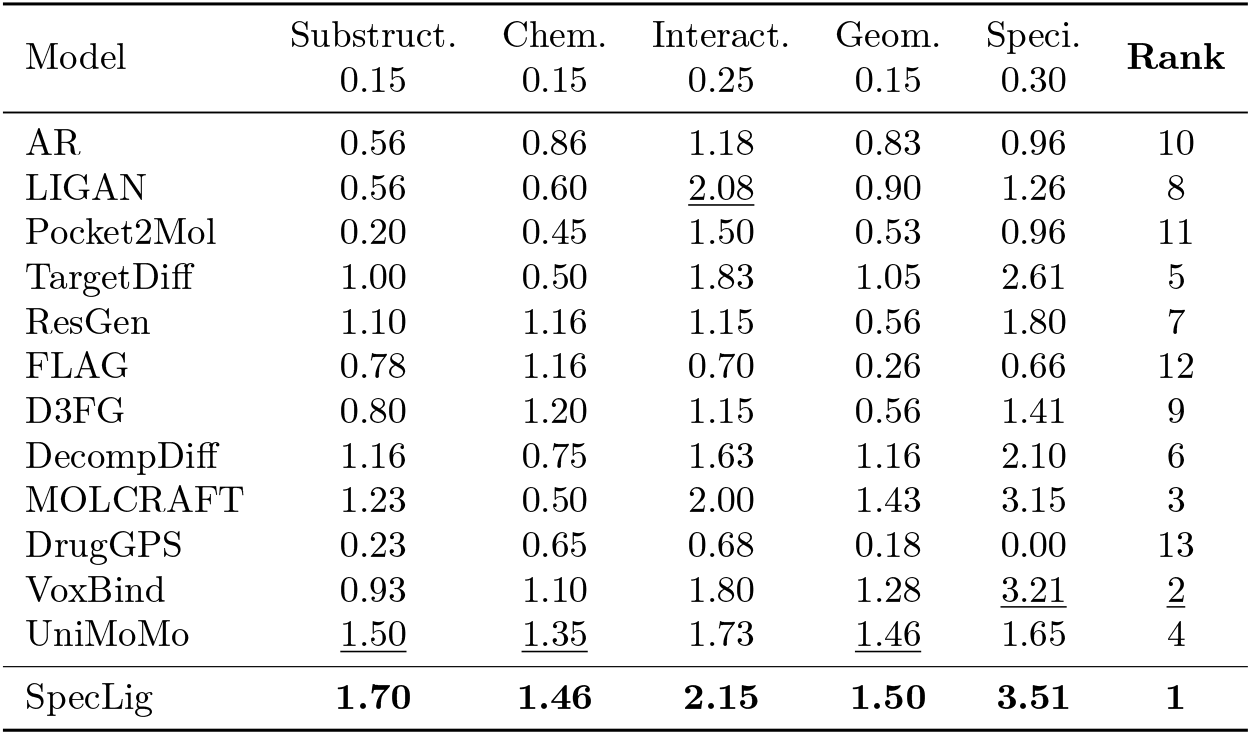
Overall comparisons for de novo small molecule design.

**Figure 2.**
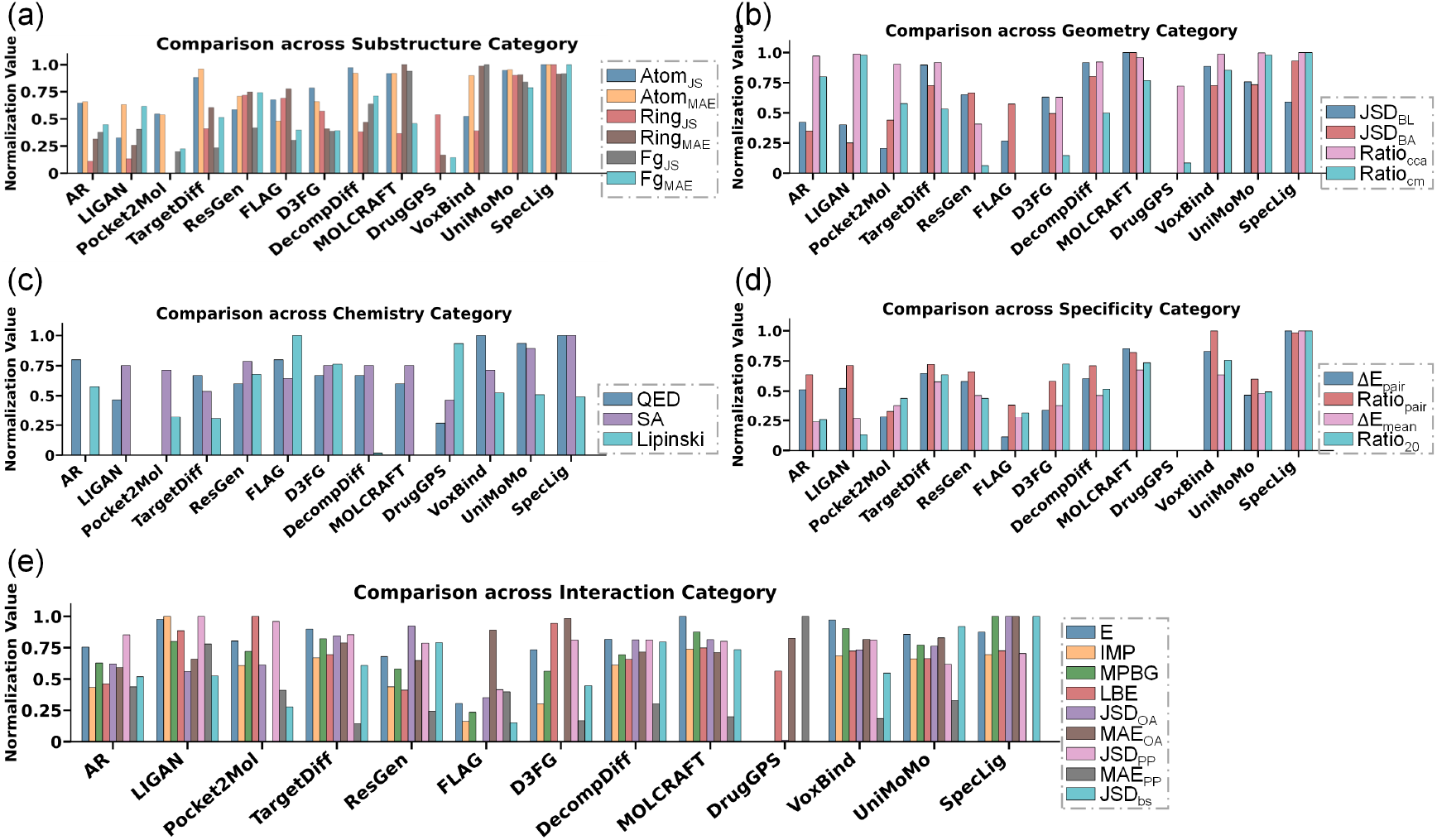
Performance comparison of small-molecule generative models across five categories of metrics: Substructure (a), Geometry (b), Chemistry (c), Specificity (d), and Interaction (e).

Despite these advances, gains in specificity and affinity are markedly smaller for small molecules than for peptides. This reflects fundamental chemical and sampling differences. The Small-molecule space is discrete and higher-dimensional: combinatorial variation in functional groups, ring systems, and rotatable bonds yields a highly multimodal energy landscape that a single block-level vocabulary sparsely covers. By contrast, amino-acid semantics are compact, so frequency-derived priors transfer more effectively. Small-molecule binding is also far more sensitive to substructure details. Therefore, overly aggressive empirical guidance induces mode collapse, while weak guidance leaves sampling inertia and reduces specificity. Thus, block–block frequency priors are a useful foundation but are insufficient alone. For small molecules, we therefore advocate integrating richer, physics-informed cues: explicit force-field or electrostatic terms, conformational-ensemble priors, fragment-aware flexible docking, adaptive guidance schedules, and targeted local geometry refinement. We believe that integrating multi-objective scoring can compensate for the limitations of frequency priors in capturing geometric details, thereby achieving a better balance between affinity and specificity.

### 3.2 Peptides

#### Setup

Referring to benchmark^43^, we organize metrics into five categories: **Specificity, Recovery, Interaction, Structural Validity**, and **Diversity. Specificity** follows the protocol described above. **Recovery** measures agree with the reference sequence and conformation. Amino-acid recovery (AAR, ↑) is the per-position sequence match rate. conformational recovery includes complex-aligned RMSD (C-RMSD, ↓) and ligand-aligned RMSD (L-RMSD, ↓). **Interaction** is primarily evaluated by the mean binding energy computed with PyRosetta (Δ*G*, ↓) and by the proportion of designed peptides that outperform the native ligand on their targets (IMP, ↑). We also report JSD_bs_, ↓ following the same definition. **Structural validity** is assessed via atomic clash rates: Clash_in_ (↓) and Clash_out_ (↓) record intra-peptide and peptide–receptor residue collisions (collision defined as *C*_*α*_ distance < 3.6574Å). Besides, backbone and side-chain dihedral deviations are summarized by JSD_bb_ (↓) and JSD_SC_ (↓) using 10^◦^ bins. **Diversity** is reported as the ratio of unique clusters to total designs (cluster threshold: sequence identity > 40% and RMSD < 2Å).

#### Baselines

Peptide baselines include RFDiffusion^23^, PepFlow^27^, PepGLAD^42^, and UniMoMo^43^. Our RFDiffusion follows the published two-stage pipeline but is limited to a single inverse-fold and relax cycle to avoid extra optimization that could bias physical metrics. PepFlow implements multimodal flow-matching for joint sequence–structure generation. PepGLAD performs latent diffusion to produce sequences and all-atom conformations. UniMoMo is the unified multimodal baseline. All methods were executed according to the recommended settings.

#### Results and discussion

Peptides, as larger ligands, are handled consistently by SpecLig. Table 2 and Figure 3 summarize SpecLig’s performance on the peptide task, and the complete results are shown in Chapter 8 of the Supplementary Information. SpecLig shows a clear advantage in the peptide task, especially in Specificity and Interaction. In Specificity (Supplementary Table S8), SpecLig increases the second-best model’s Ratiopair and Ratio20 from 68.75% and 52.91% to 75.43% and 75.00%, respectively; native ligands score 80.72% and 78.31%. In Interaction benchmarks (Supplementary Table S11), SpecLig is the only method with a negative mean Δ*G* = − 1.92, compared to 29.21 for the runner-up UniMomo. These gains reflect reduced off-target binding and higher-affinity designs. SpecLig also attains the lowest Clash_out_ and L-RMSD, indicating geometrically self-consistent outputs. A modest diversity reduction is observed, which we attribute to the constraining effect of energy guidance.

**Table 2.** Overall comparisons for de novo peptide design.

| Model | Recov.<br>0.15 | Struc.<br>0.15 | Interact.<br>0.25 | Div.<br>0.15 | Speci.<br>0.30 | Rank |
| --- | --- | --- | --- | --- | --- | --- |
| RFDiffusion | 0.00 | 0.23 | 0.00 | 0.08 | 0.15 | 5 |
| PepFlowww | 0.30 | 0.18 | 0.25 | 0.08 | 0.15 | 4 |
| PepGLAD | **0.45** | 0.18 | 0.60 | **0.60** | 0.60 | 3 |
| UniMoMo | 0.35 | **0.48** | <u>0.68</u> | <u>0.38</u> | <u>0.90</u> | <u>2</u> |
| SpecLig | <u>0.41</u> | <u>0.41</u> | **1.00** | <u>0.38</u> | **1.20** | **1** |

**Figure 3.**
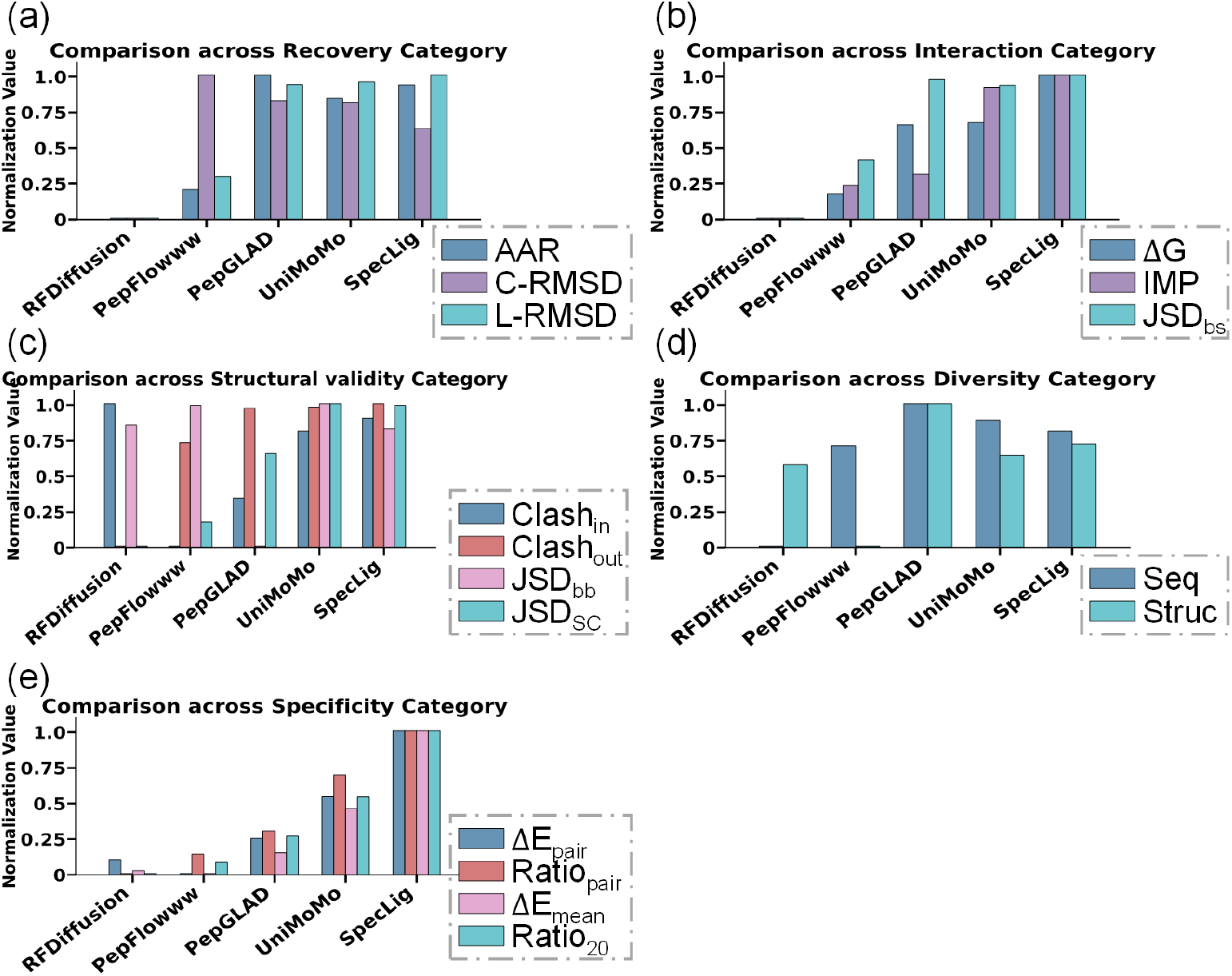
Comparison of peptide design models across five metric categories: Recovery (a), Interaction (b), Structural validity (c), Diversity (d), and Specificity (e).

Mechanistically, the gains stem from two design choices. Firstly, the hierarchical representation preserves fragment semantics and filters atom-level noise, simplifying the identification and localization of critical local units. Secondly, incorporating interaction priors into latent diffusion biases sampling toward chemically plausible, pocket-complementary solutions. Ablation studies shown in Supplementary Figure S1 confirm that removing either component degrades Specificity and Interaction. In short-peptide tasks, we observe a clear diversity–specificity tradeoff (Supplementary Figure S5): increasing energy guidance steadily reduces sequence diversity, while affinity and specificity initially rise but fall at high guidance strengths. The late decline is consistent with sampling-induced mode collapse, where over-constraining gradients concentrate probability mass on few modes and suppress beneficial exploration. In this work, we empirically tuned the guidance parameters to near-optimal values for the two design tasks. Nonetheless, SpecLig stands to benefit from more advanced schemes, such as adaptive guidance schedules implemented via gradient callbacks, entropy-aware regularization, or homology-guided template retrieval, to better reconcile specificity with diversity. Overall, SpecLig strikes a favorable balance across competing objectives, prioritizing specificity and affinity, thereby materially improving downstream candidate triage.

### 3.3 Case study

Two case studies demonstrate SpecLig’s reduction of off-target risk in small-molecule (Figure 4a–d) and peptide (Figure 4e–h) design. In both cases, SpecLig designs yielded no valid docking poses on non-target proteins (Figure 4d and 4h).

**Figure 4.**
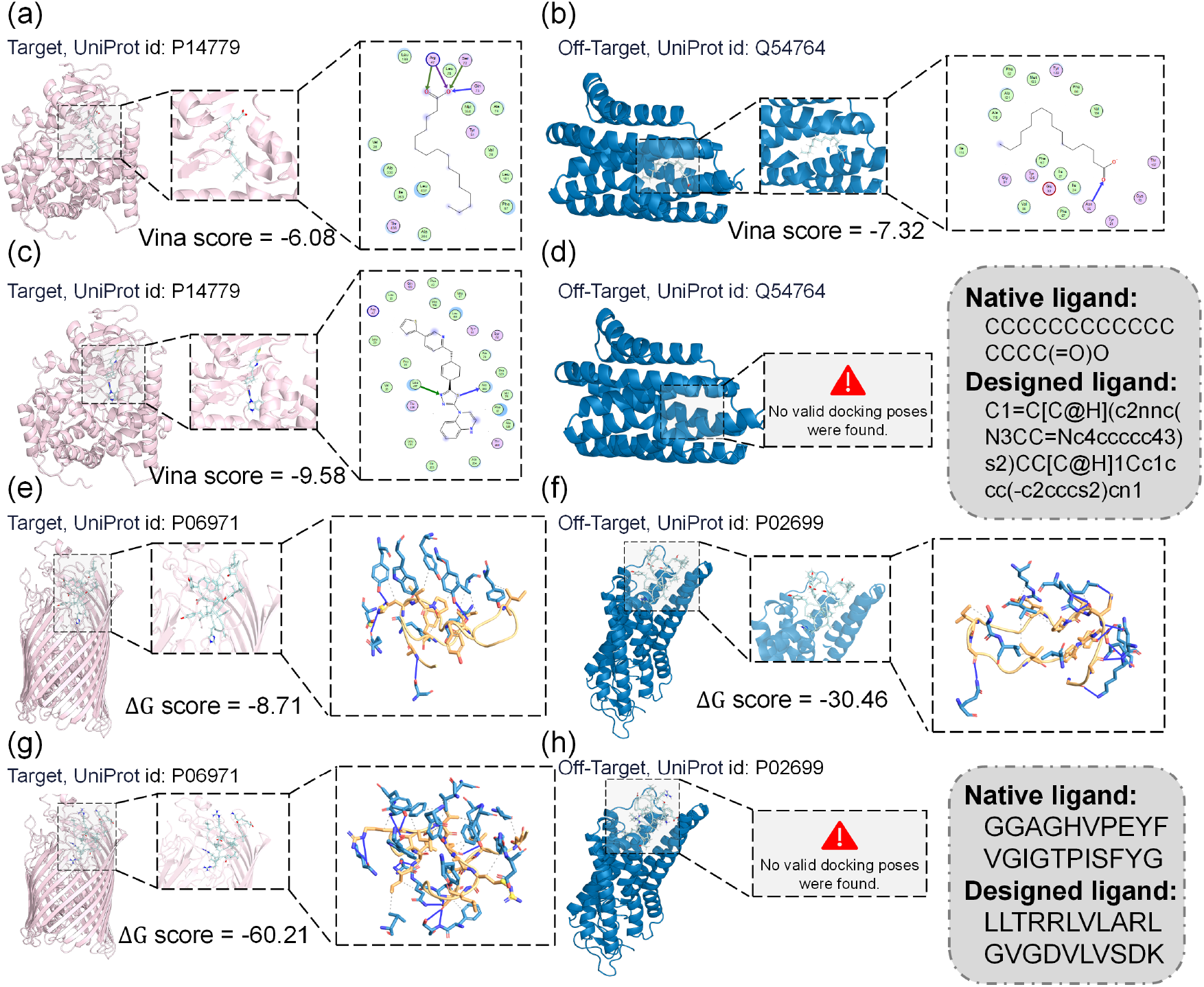
Case studies illustrating SpecLig’s reduction of off-target binding. Native small molecule targeting cytochrome P450BM-3 (a, pink) and its non-target aldehyde decarbonylase (b, blue). (c) and (d) show SpecLig-designed small molecule binds with target and non-target, separately. Similarly, (e-f) and (g-h) show the native peptide and the designed peptide binding to the target ferrichrome-iron receptor (pink) and the non-target rhodopsin (blue), respectively. All docking scores were computed using Schrödinger Glide and Rosetta under high-precision settings. Interaction analyses were performed with MOE for small molecules and the newest PLIP for peptides. The sequences of native and designed ligands are given in gray boxes.

For Cytochrome P450BM-3 (UniProt: P14779), the native ligand exhibits moderate affinity (Vina=-6.08; Figure 4a), driven by a carboxylate-mediated hydrogen-bond network with residues 47, 72, 73 and hydrophobic alkyl chain contacts. However, it binds more strongly to the non-target aldehyde decarbonylase (UniProt: Q54764; Vina=-7.32; Figure 4b) due to its topology and reusable carboxylate motif. In contrast, SpecLig’s design incorporates a thiadiazole fragment that forms directional hydrogen bonds with residues 330 and 437, meanwhile enhancing hydrophobic contacts (Vina=-9.58; Figure 4c). No candidate pose satisfied the required geometric constraints in the non-target pocket (Figure 4d), confirming SpecLig’s optimization of pocket-specific geometric and chemical complementarity.

Against the Ferrichrome-iron receptor (UniProt: P06971), native microcin J25 forms four target hydrogen bonds (Figure 4e) but nine with non-target rhodopsin (UniProt: P02699; Figure 4f), resulting in a stronger off-target affinity (-30.46) than target (-8.71). It illustrates that promiscuous contact motifs may lead to off-target risks. The SpecLig-designed peptide forms up to twelve hydrogen bonds in the target pocket, accompanied by extensive hydrophobic contacts (Figure 4g, Δ*G* = −60.21), and fails to adopt a feasible pose on the off-target (the binding energy was higher than that of the respective monomeric states, shown in Figure 4h). It demonstrates the simultaneous improvement of target affinity and suppression of non-specific binding.

Methodologically, coupling block-level modeling with an empirical energy prior shifts the model from learning generic interfacial fragments to performing pocket-conditioned fragment selection, thereby producing chemically and geometrically customized ligands. Importantly, SpecLig does not merely fill pockets by agglomerating fragments. Analyses in Supplementary Chapter 11 indicate an adaptive sizing mechanism within SpecLig: it is pocket topology, rather than blocks’ initial setting, that predominantly determines the designed ligand’s size. Lastly, additional target examples are provided in Supplementary Chapter 12.

## 4 Conclusion

SpecLig demonstrates that integrating block-wise chemical priors with hierarchical equivariant modeling yields ligand designs that better balance affinity and specificity than existing generative baselines. The hierarchical VAE reduces atom-level noise and preserves fragment semantics, while energy-guided latent sampling steers generation toward pocket-conditioned, chemically plausible solutions. Unlike black-box models, SpecLig’s preference for certain motifs can be cross-referenced with known biophysical principles, providing a rationale for the generated 3D structures. Empirical evaluation suggests consistent, interpretable improvements in specificity and interaction in both peptide and small-molecule design. Ablations verify the complementary roles of each module. Some limitations also remain. Gains in small-molecule design are constrained by discrete chemical complexity and geometric sensitivity, suggesting opportunities to incorporate richer physical cues (force-field terms, electrostatics, or conformational ensembles). Besides, prospective experimental validations are still in need. Overall, SpecLig is a pioneering step toward designing ligands with enhanced specificity and may inspire future advancements in this promising direction.

