## Supplementarty Information for "SpecLig: Energy-Guided Hierarchical Model for Target-Specific 3D Ligand Design"

### 1 Datasets and Databases

#### 1.1 CrossDock, PepBench, ProtFrag and LNR datasets

SpecLig and other structure-based small-molecule design baselines are trained and tested on CrossDock dataset, which is designed to enhance molecular design models by providing a large collection of protein-ligand complexes. This dataset specifically includes cross-docked protein-ligand pairs, which are generated by docking ligands to multiple protein targets. The primary aim is to facilitate the training and evaluation of 3D convolutional neural networks (CNNs) for predicting binding affinities and improving docking accuracy. The dataset offers a diverse set of conformations, enhancing the generalizability of models developed for drug discovery applications. The protein sequence identity between the test set and training set is constrained to be less than 30%, resulting in about 100,000 training pairs and 100 test proteins. For each pocket sample on the test set, each model generates about 100 molecules.

For peptide design, we use the training set of PepBench and PepGLAD dataset to train the models. 114 complexes of PepBench are used as validation and the LNR complexes are used as test set. PepBench is a comprehensive benchmark dataset specifically designed for protein-binding peptide design tasks. It provides curated datasets and splits of protein-peptide complexes derived from the PepGLAD framework, serving as a standardized evaluation platform for computational peptide design methods. The dataset structure includes 4,157 training complexes and 114 validation complexes, offering a substantial foundation for training and evaluating peptide design algorithms. PepBench addresses the critical need for unbiased benchmarking in the field, providing a consistent framework for comparing different methodologies.

The ProtFrag dataset represents a large-scale collection of protein fragments specifically curated for training generative models in full-atom peptide design. It contains approximately 70,498 monomer-derived protein fragments that serve as peptide-like structural building blocks for machine learning applications. The fragments in ProtFrag are extracted from protein monomers and processed to maintain structural integrity while providing the necessary diversity for model training.

The LNR (Large Non-Redundant) dataset serves as a critical test set for evaluating cross-target generalization capabilities in peptide design models. Containing 93 carefully curated non-redundant protein-peptide complexes, the LNR dataset provides a challenging benchmark that tests whether computational methods can generalize beyond their training data to novel protein targets. This dataset is particularly valuable because it consists of natural peptides that serve as high-quality references for comparing AI-generated designs. The LNR benchmark has become a standard evaluation set in recent peptide design studies.

Table S1. Sequence similarities in Datasets.

| Datasets | CrossDock | PepBench | ProtFrag | LNR |
|---|---|---|---|---|
| $Sim_{train}$ | 0.1918 | 0.2097 | 0.2544 | - |
| $Sim_{valid}$ | 0.1719 | 0.2109 | - | 0.1888 |
| $Sim_{train\text{-}valid}$ | 0.1713 | 0.1652 | - | 0.1169 |

#### 1.2 Databases

1. ChEMBL is a manually curated database of bioactive molecules with drug-like properties. It brings together chemical, bioactivity and genomic data to aid the translation of genomic information into effective new drugs. It serves as a vital resource for medicinal chemists and researchers in drug discovery, providing data for compound efficacy, safety, and pharmacological profiles. We choose all distinct compounds with complex structure in this database.
2. ZINC15 is a free database of commercially available compounds for virtual screening. It contains a vast collection of drug-like molecules, providing researchers with a resource for identifying potential drug candidates. The database is designed to facilitate the selection of compounds for experimental testing and structure-based drug design. We choose compounds that are available for purchase with LogP between -2 and 6 and molecular weight between 150 and 550.
3. PDBBind is a dataset that integrates binding affinity data with structural information from the Protein Data Bank (PDB). It features a curated collection of protein-ligand complexes, providing a valuable resource for evaluating and developing computational models for predicting binding affinities in structure-based drug design. We collect the complexes with small-molecular ligands from the in version 2020 of PDBBind.
4. Binding MOAD (Mother of All Databases) is a comprehensive, high-quality database of protein-ligand complexes with experimentally determined binding affinities. It contains a carefully curated subset of biologically relevant ligands from the Protein Data Bank, focusing on complexes with measured binding constants (Kd, Ki, IC50) and high-resolution structural data. We select complexes with binding affinity values better than 10 μM and resolution higher than 2.5 Å to ensure data quality and biological relevance for our studies.
5. PepBDB (Peptide Binding Database) is a specialized repository focusing on protein-peptide interactions, providing detailed structural and functional information about peptide binding events. It contains experimentally determined 3D structures of protein-peptide complexes, along with binding affinity data, sequence information, and functional annotations. We extract all non-redundant peptide-protein complexes with resolution better than 3.0 Å.
6. RCSB PDB (Research Collaboratory for Structural Bioinformatics Protein Data Bank) is the primary global archive for three-dimensional structural data of biological macromolecules, including proteins, nucleic acids, and their complexes. As the most comprehensive repository of macromolecular structures, it provides atomic-level insights into biological function, molecular interactions, and drug binding mechanisms. We search the entire PDB database to identify protein-protein complexes.

Table S2. Collected data in Databases.

| Database | Small molecules | Protein-small molecule complexes | Protein-protein complexes |
|---|---|---|---|
| ChEMBL | 1.7 M | - | - |
| ZINC15 | 2.8 M | - | - |
| PDBBind | - | 5 K | - |
| Binding MOAD | - | 4.5 K | - |
| PepBDB | - | - | 0.5 K |
| PDB | - | - | 30 K |

#### 2 Related Work

**Small-molecule design.** Structure-based small-molecule generation has developed into several clear paradigms. Voxel-based methods (e.g., LIGAN, VoxBind) predict spatial atomic densities and element types using variational autoencoders or walk-jump sampling on a 3D grid. SE(3)-equivariant coordinate generators (e.g., AR, Pocket2Mol, ResGen) produce atomic positions and identities directly in three-dimensional space while preserving rotational and translational invariance. Diffusion-based approaches (e.g., TargetDiff, DecompDiff) sample atomic distributions from a prior noise process and then resolve bond assignment in a post-processing step. Fragment-oriented models (FLAG, D3FG, UniMoMo) treat molecules as assemblies of rigid fragments connected by flexible bonds, substantially improving chemical realism. Strategies like MolCRAFT and DrugGPS further bias generation toward synthetically accessible regions of chemical space through continuous parameter sampling or by leveraging sub-pocket prototypes.

**Peptide design.** Peptide design has evolved from energy-based sampling toward deep generative paradigms. Flow-matching methods (e.g., PepFlow, PPFlow) extend the continuous normalizing-flow ideas to multimodal peptide representations. Geometric latent diffusion frameworks like PepGLAD combine a variational encoder with diffusion in latent space to generate peptide conformations. Separate, targeted approaches have been developed to stabilize specific peptide classes, such as α-helical peptides or D-peptides. More recently, unified frameworks (e.g., UniMoMo) have generalized peptide architectures for multi-class ligand generation and introduced iterative design schemes to improve precision.

**Block-based modeling.** Since peptides are naturally defined on amino-acid units, block-wise decomposition has been more extensively developed for small molecules. Fragment-vocabulary methods preconstruct a library of substructures and generate at the fragment level to improve synthetic accessibility. FLAG built fragments by breaking rotatable bonds and assembled 3D ligands fragment-by-fragment. UniMoMo applied a principal-subgraph extraction algorithm to derive fragments for cross-domain block graphs. With fragment libraries, retrieval-augmented or discrete flow-matching techniques (e.g., f-RAG, FragFM) have been proposed to further enhance fragment-based generation. In this work, we adopt a principal-subgraph decomposition to convert small molecules into blocks, facilitating a unified representation compatible with the amino-acid–level peptide modeling.

**Energy-guided models.** In protein design, diffusion generators increasingly incorporate empirical potentials or custom energy terms during or after sampling to refine outputs (e.g., RFdiffusion, Chroma). TCRen used statistical and energy priors to guide the generation of T-cell receptor. In small-molecule design, customized energy terms have been proposed to optimize specified properties. Gao et al. applied contrastive learning and energy terms to guide generative orientation, partially alleviating the off-target problem. DiffSBDD demonstrated the application of additional constraints to improve the generated candidate drugs based on expected computational metrics. Subsequent methods like ALIDIFF and IPDIFF introduced distribution alignment and interaction-prior methods to inject interaction-aware guidance into the diffusion process.

### 3 Experimental Details about Specificity Analysis

To systematically evaluate the target specificity of generative models in structure-based drug design (SBDD), we established a comprehensive computational framework. For each designed ligand (both small molecules and peptides), we performed molecular docking against the intended target protein and 100 randomly selected off-target proteins from the PDB database. The off-target selection was constrained to exclude homologous proteins (sequence identity >30%). AutoDock Vina and Rosetta were used for docking. We defined ligand specificity based on comparative binding affinity analysis. A ligand was classified as "target-specific" if its calculated binding affinity ($\Delta G$) for the intended target was stronger than 90% of its affinities for the 100 off-target proteins. Conversely, ligands whose target affinity ranked below the 90% off-target affinities were categorized as "non-specific."

For small molecule ligands, we quantified the fraction of polar functional groups (shown in Table S3) relative to total heavy atoms. This metric was calculated using RDKit's chemical feature perception algorithms with standard SMILES parsing. For peptide ligands, secondary structure composition was determined using DSSP algorithm applied to the lowest-energy docking poses, with helical content defined as the percentage of residues adopting α-helical conformations.

Across 12 advanced small-molecule generative models, non-specific ligands exhibited a significantly higher proportion of polar groups (mean = 19.9% ± 7.9%) compared to their specific counterparts (mean = 25.4% ± 6.2%). This elevated polar group density facilitates non-selective hydrogen bonding networks across diverse protein surfaces, explaining the observed promiscuity. For the peptide ligands designed by 4 advanced models, helical content emerged as a critical determinant of specificity. Target-specific peptides typically maintained a higher fraction of helical conformations, about 3%-10%. This structural preference arises because well-defined helical conformations reduce entropic penalties upon binding and minimize solvent-exposed flexible segments that could engage in non-specific interactions. These findings indicate a common tension in generative SBDD models: while model architectures naturally favor frequently occurring chemical motifs and secondary structures, these features do not inherently optimize target discrimination.

Table S3. Polar groups defined for statistics.

| Group name | SMILES |
|---|---|
| Alcohol hydroxyl | [OX2H] |
| phenol | c[OX2H] |
| Carboxylic acid | C(=O)[OX2H1] |
| ester | C(=O)O[!#1] |
| amide | C(=O)N |
| Primary amine | [NX3;H2] |
| Secondary amine | [NX3;H1] |
| Tertiary amine | [NX3;H0] |
| nitro | [NX3](=O)=O |
| thiol | [SX2H] |
| sulfonyl | S(=O)(=O)[#6] |
| ether | O([#6])[#6] |

### 4 Algorithms for Training with SpecLig

We provide the training pseudocode of SpecLig below, including two steps for training hierarchical VAE and LDM.

**Algorithm 1** Training Algorithm for SpecLig

**Input**: Complex dataset $D = \{(G_P, G_L)\}$, block vocabulary $S$
**Output**: Trained encoder $\mathcal{E}_\xi = \{\mathcal{E}_{\xi,1}, \mathcal{E}_{\xi,2}\}$, decoder $\mathcal{D}_\phi = \{\mathcal{D}_{\phi,1}, \mathcal{D}_{\phi,2}\}$, and latent diffusion model $\boldsymbol{\varepsilon}_\theta$

**Stage I: Train hierarchical VAE**
**Function** $\text{Encode}(G, \mathcal{E}_\xi)$
  $\{(\boldsymbol{\mu}_i, \boldsymbol{\sigma}_i, \vec{\boldsymbol{\mu}}_i, \vec{\boldsymbol{\sigma}}_i)\} \leftarrow \mathcal{E}_\xi(G)$ // get the latent distribution of representations
  Sample $\{(\varepsilon, \vec{\varepsilon})\} \sim \mathcal{N}(0, \boldsymbol{I})$
  $\boldsymbol{z}_i \leftarrow \boldsymbol{\mu}_i + \boldsymbol{\sigma}_i \odot \varepsilon,\ \vec{\boldsymbol{z}}_i \leftarrow \vec{\boldsymbol{\mu}}_i + \vec{\boldsymbol{\sigma}}_i \odot \varepsilon$ // reparameterized sampling
  $Z \leftarrow \{(\boldsymbol{z}_i, \vec{\boldsymbol{z}}_i)\}$
  **return** $Z$
**end Function**
Initialize $\mathcal{E}_\xi, \mathcal{D}_\phi$
**while** $\xi, \phi$ have not converged **do**
  Randomly sample $(G_P, G_L) \sim D$
  $Z_P^0, Z_L^0 \leftarrow \text{Encode}(G_P, \mathcal{E}_\xi), \text{Encode}(G_L, \mathcal{E}_\xi)$
  $Z_L^0 \leftarrow Z_L^0 + \omega_{inj} \cdot \mathcal{N}(0, \boldsymbol{I})$ // inject gaussian noise for robustness
  Randomly sample indices $\tilde{\mathbb{I}}_P \subseteq \mathbb{I}_P$ for 5% of pocket residues
  $\mathbb{I} \leftarrow \tilde{\mathbb{I}}_P \cup \mathbb{I}_L$
  $\{(h_i, \vec{r}_i | i \in \mathbb{I})\} \leftarrow \mathcal{D}_{\phi,2}(Z_P^0, Z_L^0)$ // get the block-scale attribute and position latents
  $\hat{s}_i \leftarrow \text{MLP}_{block}(h_i)$
  $(\boldsymbol{\alpha}_i, \vec{\boldsymbol{x}}_i, \boldsymbol{\beta}_i) = \text{Lookup}(\sigma(s_i), S)$ // get the element types and intra-block bonds
  Initialize $\vec{\boldsymbol{x}}_i^1 \sim \mathcal{N}(\vec{r}_i, \boldsymbol{I}), t \sim U(0,1)$
  $\vec{\boldsymbol{x}}_i^t \leftarrow t \cdot \vec{\boldsymbol{x}}_i^1 + (1-t) \cdot \vec{\boldsymbol{x}}_i^0$ // get the noisy position from ground truth and prediction
  $\overline{\boldsymbol{\beta}}_{ij,ab} \leftarrow \emptyset$ if $p < 0.5$ else $\boldsymbol{\beta}_{ij,ab}, p \sim U(0,1)$ // provide 50% inter-block bonds
  $\{h_i, \Delta\vec{\boldsymbol{x}}_i | i \in \mathbb{I}\} \leftarrow \mathcal{D}_{\phi,1}(\{(\boldsymbol{\alpha}_i, \vec{\boldsymbol{x}}_i^t, \boldsymbol{\beta}_i \cup \overline{\boldsymbol{\beta}}_{ij,ab})\}, Z_P^0, Z_L^0, G_P, t)$ // get the atom-scale latents
  $p(a,b) \leftarrow \text{MLP}_{focal}\left(h_i^0, h_j^0, d(\vec{\boldsymbol{x}}_i^0, \vec{\boldsymbol{x}}_j^0)\right)$ // compute frontier scores of atom-pair
  $\hat{\boldsymbol{\beta}}_{ij,ab} \leftarrow \text{MLP}_{bond}(h_{i,a}^0, h_{j,b}^0)$ // predict the bond types
  $\mathcal{L}_{VAE} \leftarrow \sum_{i\in\mathbb{I}} \left(\mathcal{L}_{VAE-atom}(i) + \mathcal{L}_{VAE-block}(i) + \mathcal{L}_{VAE-repr}(i)\right) / |\mathbb{I}|$
  $\xi, \phi \leftarrow \text{optimizer}(\mathcal{L}_{VAE}; \xi, \phi)$
**end while**
**return** $\mathcal{E}_\xi,\ \mathcal{D}_\phi$

**Stage II: Train latent diffusion model**
Fix $\mathcal{E}_\xi,\ \mathcal{D}_\phi$, initialize $\boldsymbol{\varepsilon}_\theta$.
**while** $\theta$ has not converged **do**
  Randomly sample $(G_P, G_L) \sim D$
  $Z_P^0, Z_L^0 \leftarrow \text{Encode}(G_P, \mathcal{E}_\xi), \text{Encode}(G_L, \mathcal{E}_\xi)$
  Initialize $t \sim U(0,1), \{(\varepsilon, \vec{\varepsilon})\} \sim \mathcal{N}(0, \boldsymbol{I})$
  $Z_L^t \leftarrow \{(\boldsymbol{z}_i^t, \vec{\boldsymbol{z}}_i^t) | i \in \mathbb{I}_L, [\boldsymbol{z}_i^t, \vec{\boldsymbol{z}}_i^t] = \sqrt{\bar{\alpha}_t}[\boldsymbol{z}_i^0, \vec{\boldsymbol{z}}_i^0] + \sqrt{1-\bar{\alpha}_t}[\varepsilon_i, \vec{\varepsilon}_i]\}$
  $\hat{\boldsymbol{\varepsilon}} \leftarrow \boldsymbol{\varepsilon}_\theta(Z_L^t, Z_P^0, t)$
  $\hat{Z}_L^0 \leftarrow \text{rev}(Z_L^t, \hat{\boldsymbol{\varepsilon}})$ // get the "clean" sample form predicted noise
  $\mathcal{L}_{LDM} \leftarrow \sum_{i\in\mathbb{I}} \left(\|\hat{\boldsymbol{\varepsilon}} - \{(\varepsilon, \vec{\varepsilon})\}\|^2 + \mathcal{L}_{LPL}(\mathcal{D}_{\phi,2}, \hat{Z}_L^0, Z_L^0)\right) / |\mathbb{I}|$
  $\theta \leftarrow \text{optimizer}(\mathcal{L}_{LDM}; \theta)$
**end while**
**return** $\boldsymbol{\varepsilon}_\theta$

### 5 Algorithms for Sampling with SpecLig

We provide the sampling pseudocode of SpecLig below, energy-guided LDM is the core module of sampling.

**Algorithm 2** Energy-guided Sampling for SpecLig

**Input**: Pocket $G_P$, encoder $\mathcal{E}_\xi$, decoder $\mathcal{D}_\phi$, latent diffusion model $\boldsymbol{\varepsilon}_\theta$, block vocabulary $S$, and block-wise prior $F$

**Output**: Generated ligand $G_L$

**Function** $\mathrm{Decode}\big(G_P, \mathcal{D}_{\phi,1}, \{(\boldsymbol{\alpha_i}, \overline{\boldsymbol{\beta}}_{ij,ab}, \boldsymbol{\beta_i})\}, Z_P^0, Z_L^0\big)$
    Initialize $\vec{\boldsymbol{x}}_i^1 \sim \mathcal{N}(\vec{r}_i, \boldsymbol{I})$
    **for** $t = 1.0$ **to** 0.0 step $-\Delta t$ **do**
        $h_i^t, \Delta\vec{\boldsymbol{x}}_i^t \leftarrow \mathcal{D}_{\phi,1}\big(\{(\boldsymbol{\alpha_i}, \vec{\boldsymbol{x}}_i^t, \boldsymbol{\beta_i} \cup \overline{\boldsymbol{\beta}}_{ij,ab})\}, Z_P^0, Z_L^0, G_P, t\big)$ // predict the adjustment in position
        $h_i^{t-\Delta t} \leftarrow h_i^t$
        $\vec{\boldsymbol{x}}_i^{t-\Delta t} \leftarrow \vec{\boldsymbol{x}}_i^t + \Delta\vec{\boldsymbol{x}}_i^t \cdot \Delta t$
    **end for**
    $p(a,b) \leftarrow \mathrm{MLP}_{focal}\left(h_i^0, h_j^0, d(\vec{\boldsymbol{x}}_i^0, \vec{\boldsymbol{x}}_j^0)\right)$
    $\boldsymbol{\beta}_{ij} \leftarrow \left\{\sigma\left(\mathrm{MLP}_{bond}(h_{i,a}^0, h_{j,b}^0)\right) \middle| i \neq j, dist^0 < 3.5\text{Å}, p(a,b) > thre\right\}$ if $\boldsymbol{\beta}_{ij} = \emptyset$ else $\boldsymbol{\beta}_{ij}$
    **return** $\{\vec{\boldsymbol{x}}_i^t, \boldsymbol{\beta}_{ij}\}$
**end Function**
Initialize $Z_L^T \sim \mathcal{N}(0, \boldsymbol{I})$
$Z_P^0 \leftarrow \mathrm{Encode}(G_L, \mathcal{E}_\xi)$
**for** $t$ in $T, T-1, \dots, 1$ **do**
    $\hat{\boldsymbol{\varepsilon}} \leftarrow \boldsymbol{\varepsilon}_\theta(Z_L^t, Z_P^0, t)$
    $\hat{Z}_L^0 \leftarrow \mathrm{rev}(Z_L^t, \hat{\boldsymbol{\varepsilon}})$ // get the "clean" sample form predicted noise
    $\hat{\mathrm{s}} \leftarrow \mathrm{MLP}_{block}\left(\mathcal{D}_{\phi,2}(\hat{Z}_L^0)\right)$ // get the potential distribution of block type
    $\mathrm{s} \leftarrow \hat{\mathrm{s}} \cup \mathrm{s_P}$ // include the block type from ground truth
    $E \leftarrow \sum_{i,j} -\log\left(\left(s_i{}^T F s_j\right)^{1/\tau}\right) \cdot \omega_{ij}$
    $\hat{\boldsymbol{\varepsilon}}_{guide} \leftarrow \hat{\boldsymbol{\varepsilon}} - \omega_t \cdot \mathrm{ClipNorm}(\nabla_{\hat{\boldsymbol{\varepsilon}}} E)$ // guide the predicted noise
    $\hat{Z}_L^{t-1} \leftarrow \frac{1}{\sqrt{\alpha_t}}\left(\hat{Z}_L^t - \frac{\beta_t}{\sqrt{1-\bar{\alpha}_t}}\hat{\boldsymbol{\varepsilon}}_{guide}\right) + \beta_t \varepsilon$
**end for**
$\{(h_i, \vec{r}_i | i \in \mathbb{I}_L)\} \leftarrow \mathcal{D}_{\phi,2}(Z_P^0, \hat{Z}_L^0)$
$\hat{\mathrm{s}}_i \leftarrow \mathrm{MLP}_{block}(h_i)$
$(\boldsymbol{\alpha_i}, \vec{\boldsymbol{x}}_i, \boldsymbol{\beta_i}) = \mathrm{Lookup}(\sigma(\hat{\mathrm{s}}_i), S)$
$\{(\vec{\boldsymbol{x}}_i^0, \boldsymbol{\beta}_{ij}) | i \in \mathbb{I}_L, j \in \mathbb{I}\} \leftarrow \mathrm{Decode}\big(G_P, \mathcal{D}_{\phi,1}, \{(\boldsymbol{\alpha_i}, \emptyset, \boldsymbol{\beta_i})\}, Z_P^0, Z_L^0\big)$
$G_L \leftarrow \{(\boldsymbol{\alpha_i}, \vec{\boldsymbol{x}}_i^0, \boldsymbol{\beta_i} \cup \boldsymbol{\beta}_{ij})\}$
$Z_L \leftarrow \mathrm{Encode}(G_L, \mathcal{E}_\xi)$ // Re-encode for refinement
$\{(\vec{\boldsymbol{x}}_i^0, \boldsymbol{\beta}_{ij}) | i \in \mathbb{I}_L, j \in \mathbb{I}\} \leftarrow \mathrm{Decode}\big(G_P, \mathcal{D}_{\phi,1}, \{(\boldsymbol{\alpha_i}, \boldsymbol{\beta}_{ij}, \boldsymbol{\beta_i})\}, Z_P^0, Z_L^0\big)$ // Refinement with inter-block bonds
$G_L \leftarrow \{(\boldsymbol{\alpha_i}, \vec{\boldsymbol{x}}_i^0, \boldsymbol{\beta_i} \cup \boldsymbol{\beta}_{ij})\}$
**return** $G_L$

### 6 Latent Perceptual Loss

While latent perceptual loss (LPL) has demonstrated effectiveness in image generation tasks, its direct application to 3D molecular generation presents unique challenges due to the discrete nature of chemical structures and the critical importance of geometric fidelity. We adapt LPL to the ligand design domain by leveraging the hierarchical decoder's intermediate representations to preserve structural plausibility during denoising process.

Specifically, given a noisy latent representation $\boldsymbol{z}_t$ at diffusion step $t$, our denoising network $\varepsilon_\theta$ predicts noise $\hat{\varepsilon}$, from which we could reconstruct the clean estimate $\hat{\boldsymbol{z}} = \left(\boldsymbol{z}_t - \sqrt{1-\bar{\alpha}_t}\hat{\varepsilon}\right)/\sqrt{\bar{\alpha}_t}$. The LPL loss computes the perceptual difference between features extracted from decoding $\hat{\boldsymbol{z}}$ and the ground-truth latent $\boldsymbol{z}_0$:

$$\mathcal{L}_{LPL} = \sum_{l=1}^{L} \omega_l \left\| \pi_l\left(\mathcal{D}_{\phi,2}(\hat{\boldsymbol{z}}, Z_P)\right) - \pi_l\left(\mathcal{D}_{\phi,2}(\boldsymbol{z}_0, Z_P)\right) \right\|_2 \quad (1)$$

Where $\pi_l(\cdot)$ denotes the feature map at layer $l$ of our block-scale decoder $\mathcal{D}_{\phi,2}$, $Z_P$ is the encoded representation of pocket, and $\omega_l$ balances contributions across different layers. Crucially, we restrict LPL application to fine-tuning stages and only when the signal-to-noise ratio exceeds $\tau_{snr} = 0.6$, as early diffusion steps lack sufficient structural information for meaningful perceptual comparison. We further enhance training stability by clipping feature differences exceeding three standard deviations from their rolling mean, preventing outlier gradients from destabilizing optimization. LPL loss partially bridges the gap between VAE and latent diffusion models, ultimately yielding ligands with improved synthetic accessibility and drug-likeness.

### 7 Full Numerical Results for the Small Molecule Design

#### 7.1 Specificity related metrics

We performed two complementary cross-docking validation experiments to quantify off-target binding, and peptides and small molecules were treated equivalently. In Experiment 1, each binder was docked to its target pocket and to one randomly selected non-target pocket using the same high-precision protocol; this produces a paired target vs. single off-target energy for each binder. In Experiment 2, we selected a random subset of designed binders (e.g., M = 20) and docked each subset binder to every pocket in the full dataset using a faster, lower-resolution but computationally cheap docking model. It produces a full ranking of the target pocket among all pockets for each binder.

**$\Delta E_{pair}$ (↓)** means pair wise energy difference: for each binder in Experiment 1 we compute $\Delta E_{pair} = E_{target} - E_{cross}$, negative values indicate the binder is predicted to bind the target better than the random sampled pocket. **$Ratio_{pair}$ (↑)** calculates the proportion of binders in Experiment 1 for which $E_{target} < E_{cross}$. **$\Delta E_{mean}$ (↓)** calculates the mean energy advantage over all pockets in Experiment 2, it is computed by $E_{target} - \overline{E}_{all}$. **$Ratio_{20}$ (↑)** reports the fraction of binders whose target energy rank falls within the best 20% in Experiment 2. Then we could get the comparison results below.

Table S4. Complete comparison results for small molecule design under Specificity category

| Method | $\Delta E_{pair}$ | $Ratio_{pair}$ | $\Delta E_{mean}$ | $Ratio_{20}$ | **Rank** |
|---|---|---|---|---|---|
| Reference | -1.18 | 58.60 | -0.81 | 38.78 | - |
| AR | -0.40 | 51.96 | -0.21 | 21.75 | 9.8 |
| LIGAN | -0.41 | 53.44 | -0.23 | 20.33 | 8.8 |
| Pocket2Mol | -0.20 | 46.03 | -0.31 | 23.82 | 9.8 |
| TargetDiff | -0.52 | 53.65 | -0.45 | 26.03 | 4.3 |
| ResGen | -0.46 | 52.44 | -0.37 | 23.82 | 7.0 |
| FLAG | -0.05 | 47.07 | -0.24 | 22.45 | 10.8 |
| D3FG | -0.25 | 50.89 | -0.31 | 27.06 | 8.3 |
| DecompDiff | -0.48 | 53.41 | -0.37 | 24.70 | 6.0 |
| MOLCRAFT | <u>-0.70</u> | 55.55 | <u>-0.52</u> | 27.17 | 2.5 |
| DrugGPS | 0.05 | 39.59 | -0.04 | 18.84 | 13.0 |
| VoxBind | -0.68 | **59.07** | -0.49 | <u>27.41</u> | <u>2.3</u> |
| UniMoMo | -0.36 | 51.25 | -0.38 | 24.44 | 7.5 |
| SpecLig | **-0.83** | <u>58.73</u> | **-0.75** | **30.17** | **1.3** |

#### 7.2 Chemistry related metrics

The evaluation of chemical properties builds on prior work, incorporating several key metrics. These include **QED (↑)**, which provides a quantitative estimation of drug-likeness; **SA (↑)**, the synthetic accessibility score normalized to [0, 1], with higher scores indicating better synthesizability; and **LogP** (-), the octanol-water partition coefficient, with optimal values typically ranging between -0.4 and 5.6 for drug candidates. Additionally, the **LPSK (↑)** metric measures the proportion of generated drug molecules that satisfy Lipinski's rule of five, a guideline for assessing drug-like properties.

Table S5. Complete comparison results for small molecule design under Chemistry category

| Method | QED | LogP | SA | Lipinski | **Rank** |
|---|---|---|---|---|---|
| AR | 0.51 | 0.63 | 0.45 | 4.73 | 7.3 |
| LIGAN | 0.46 | 0.56 | 0.66 | 4.39 | 9.0 |
| Pocket2Mol | 0.39 | 2.39 | 0.65 | 4.58 | 10.0 |
| TargetDiff | 0.49 | 1.13 | 0.60 | 4.57 | 9.7 |
| ResGen | 0.48 | 1.61 | 0.67 | 4.79 | 5.3 |
| FLAG | 0.51 | 1.62 | 0.63 | **4.98** | 5.3 |
| D3FG | 0.49 | 1.56 | 0.66 | 4.84 | 5.0 |
| DecompDiff | 0.49 | 1.22 | 0.66 | 4.40 | 8.0 |
| MOLCRAFT | 0.48 | 0.87 | 0.66 | 4.39 | 9.7 |
| DrugGPS | 0.43 | 0.12 | 0.58 | <u>4.94</u> | 8.7 |
| VoxBind | **0.54** | 2.22 | 0.65 | 4.70 | 5.7 |
| UniMoMo | <u>0.53</u> | 1.51 | <u>0.7</u> | 4.69 | <u>4.0</u> |
| SpecLig | **0.54** | 1.46 | **0.73** | 4.68 | **3.3** |

#### 7.3 Interaction related metrics

The evaluation of interactions encompasses two primary aspects: Vina scores and interaction distributions. **E (↓)** represents the average Vina energy, while **IMP(%) (↑)** calculates the percentage of generated molecules with better Vina energy than the reference molecules.

Two metrics are further introduced to reduce the bias of Vina energy on the size of molecules: mean percent binding gap (MPBG) and ligand binding efficiency (LBE). **MPBG (↑)** measures relative improvement of generated molecules over reference molecules in terms of Vina energy for each binding pocket. **LBE (↑)** evaluates the average energy contribution per atom.

Interaction distribution metrics assess the similarity between generated and reference molecules across seven interaction types identified by PLIP software. **$JSD_{OA}$ (↓)** and **$MAE_{OA}$ (↓)** evaluate overall distributions of interaction types, while **$JSD_{PP}$ (↓)** and **$MAE_{PP}$ (↓)** focus on per-pocket distributions. Besides, we denote **$JSD_{bs}$ (↓)** as the Jensen–Shannon distance between the block distribution of the designed ligand and the block distribution of the reference ligand within the binding site.

Table S6. Complete comparison results for small molecule design under Interaction category

| Method | E | IMP% | MPBG% | LBE | $JSD_{OA}$ | $MAE_{OA}$ | $JSD_{PP}$ | $MAE_{PP}$ | $JSD_{bs}$ | **Rank** |
|---|---|---|---|---|---|---|---|---|---|---|
| AR | -6.86 | 37.18 | -5.28 | 0.3093 | 0.0314 | 0.1060 | 0.1759 | 0.4097 | 0.1611 | 8.3 |
| LIGAN | <u>-7.70</u> | **72.71** | 4.22 | 0.3897 | 0.0346 | 0.0905 | **0.1451** | <u>0.3416</u> | 0.1603 | <u>4.7</u> |
| Pocket2Mol | -7.05 | 48.07 | -0.17 | **0.4115** | 0.0319 | 0.2455 | <u>0.1535</u> | 0.4152 | 0.2067 | 7.0 |
| TargetDiff | -7.41 | 51.99 | 5.38 | 0.3537 | 0.0198 | 0.0600 | 0.1757 | 0.4687 | 0.1441 | 5.7 |
| ResGen | -6.58 | 37.47 | -7.97 | 0.2997 | <u>0.0157</u> | 0.0926 | 0.1902 | 0.4487 | 0.1098 | 8.4 |
| FLAG | -5.17 | 20.00 | -26.88 | 0.2215 | 0.0455 | 0.0360 | 0.2684 | 0.4179 | 0.2307 | 10.2 |
| D3FG | -6.78 | 28.90 | -8.85 | <u>0.4009</u> | 0.0638 | <u>0.0135</u> | 0.1850 | 0.4641 | 0.1750 | 8.4 |
| DecompDiff | -7.10 | 48.31 | -1.59 | 0.3460 | 0.0215 | 0.0769 | 0.1848 | 0.4369 | 0.1085 | 6.5 |
| MOLCRAFT | **-7.79** | <u>56.22</u> | 8.38 | 0.3638 | 0.0214 | 0.0780 | 0.1868 | 0.4574 | 0.1205 | 5.0 |
| DrugGPS | -4.02 | 9.99 | -39.74 | 0.3287 | 0.0632 | 0.0509 | 0.3555 | **0.2973** | 0.2593 | 10.3 |

| | | | | | | | | | | |
|---|---|---|---|---|---|---|---|---|---|---|
| VoxBind | -7.68 | 52.91 | 9.89 | 0.3588 | 0.0257 | 0.0533 | 0.1850 | 0.4606 | 0.1557 | 5.8 |
| UniMoMo | -7.25 | 51.4 | 2.63 | 0.3473 | 0.0241 | 0.0503 | 0.2256 | 0.4316 | 0.0854 | 6.1 |
| SpecLig | -7.32 | 53.6 | **15.17** | 0.3593 | **0.0117** | **0.0099** | 0.2074 | 0.4973 | **0.0700** | **4.4** |

#### 7.4 Substructure related metrics

The comparison focuses on atom types, ring types, and functional groups, assessing how closely the generated distributions align with the reference. Two types of metrics are involved: Jensen-Shannon divergence (JSD), and mean absolute error (MAE). **JSD (↓)** calculates the divergence between the overall probablistic distributions on the specific substructures of the generated and the reference molecules. **MAE (↓)** calculates the difference between the molecule-level occurring frequencies of substructures (e.g. carbon atoms occur 16 times in one molecule on average) of the generated and reference molecules.

Table S7. Complete comparison results for small molecule design under Substructure category

| | $\text{Atom}_{JS}$ | $\text{Atom}_{MAE}$ | $\text{Ring}_{JS}$ | $\text{Ring}_{MAE}$ | $\text{Fg}_{JS}$ | $\text{Fg}_{MAE}$ | **Rank** |
|---|---|---|---|---|---|---|---|
| AR | 0.0803 | 0.8183 | 0.3227 | 0.2514 | 0.2536 | 0.0482 | 9.3 |
| LIGAN | 0.1167 | 0.8680 | 0.3163 | 0.2701 | 0.2468 | 0.0378 | 9.3 |
| Pocket2Mol | 0.0916 | 1.0497 | 0.3550 | 0.3545 | 0.2961 | 0.0622 | 11.7 |
| TargetDiff | 0.0533 | 0.2399 | 0.2345 | 0.1559 | 0.2876 | 0.0441 | 6.3 |
| ResGen | 0.0872 | 0.7245 | 0.1436 | 0.1086 | 0.2438 | 0.0299 | 5.7 |
| FLAG | 0.0768 | 1.1598 | 0.1517 | 0.0992 | 0.2706 | 0.0512 | 7.8 |
| D3FG | 0.0644 | 0.8154 | 0.1869 | 0.2204 | 0.2511 | 0.0516 | 7.7 |
| DecompDiff | 0.0431 | 0.3197 | 0.2431 | 0.2006 | 0.1916 | 0.0318 | 5.3 |
| MOLCRAFT | 0.0490 | 0.3208 | 0.2469 | **0.0264** | 0.1196 | 0.0477 | 4.8 |
| DrugGPS | 0.1539 | 2.0802 | 0.1964 | 0.2998 | 0.3433 | 0.0671 | 11.5 |
| VoxBind | 0.0942 | 0.3564 | 0.2401 | 0.0301 | **0.1053** | 0.0761 | 6.8 |
| UniMoMo | 0.0458 | 0.2559 | 0.0897 | 0.0573 | 0.1433 | 0.0271 | 3.0 |
| SpecLig | **0.0398** | **0.1678** | **0.0613** | 0.0548 | 0.1251 | **0.0139** | **1.7** |

#### 7.5 Geometry related metrics

This group assesses the authenticity of generated molecules by analyzing local geometries, including bond lengths, bond angles, and atomic clashes. $\mathbf{JSD_{BL}}$ (↓) quantifies the Jensen-Shannon divergence between the bond length distributions of generated and reference molecules. Similarly, $\mathbf{JSD_{BA}}$ (↓) evaluates the divergence for bond angles, which are discretized every two degrees between 0° to 180°. Clashes between generated molecules and target proteins are identified based on van der Waals radius overlaps of ⩾ 0.4Å. $\mathbf{Ratio_{cca}}$ (↓) measures the average proportion of clashing atoms, while $\mathbf{Ratio_{cm}}$ (↓) reflects the average proportion of molecules with at least one clashing atom.

Table S8. Complete comparison results for small molecule design under Geometry category

| Method | $\text{JSD}_{BL}$ | $\text{JSD}_{BA}$ | $\text{Ratio}_{cca}$ | $\text{Ratio}_{cm}$ | **Rank** |
|---|---|---|---|---|---|
| AR | 0.4564 | 0.5275 | 0.0190 | 0.2395 | 7.5 |
| LIGAN | 0.4645 | 0.5673 | 0.0096 | 0.0718 | 7.0 |
| Pocket2Mol | 0.5433 | 0.4922 | 0.0576 | 0.4499 | 9.5 |

| | | | | | |
|---|---|---|---|---|---|
| TargetDiff | 0.2659 | 0.3769 | 0.0483 | 0.4920 | 6.0 |
| ResGen | 0.3641 | 0.4020 | 0.3368 | 0.9331 | 9.3 |
| FLAG | 0.5184 | 0.4385 | 0.5667 | 0.9930 | 11.3 |
| D3FG | 0.3727 | 0.4700 | 0.2115 | 0.8571 | 9.3 |
| DecompDiff | <u>0.2576</u> | 0.3473 | 0.0462 | 0.5248 | 5.3 |
| MOLCRAFT | **0.2250** | **0.2683** | 0.0264 | 0.2691 | 3.5 |
| DrugGPS | 0.6249 | 0.6672 | 0.1596 | 0.9133 | 11.8 |
| VoxBind | 0.2701 | 0.3771 | 0.0103 | 0.1890 | 4.5 |
| UniMoMo | 0.3217 | 0.3746 | <u>0.0041</u> | <u>0.0714</u> | <u>3.3</u> |
| SpecLig | 0.3886 | <u>0.2962</u> | **0.0030** | **0.0505** | **3.0** |

### 8 Full Numerical Results for the Peptide Design

#### 8.1 Specificity related metrics

For Specificity related metrics, we follow the same definition in small molecule design.

Table S9. Complete comparison results for peptide design under Specificity category

| Method | $\Delta E_{pair}$ | $Ratio_{pair}$ | $\Delta E_{mean}$ | $Ratio_{20}$ | **Rank** |
|---|---|---|---|---|---|
| Reference | -25.29 | 80.72% | -26.41 | 78.31% | - |
| RFDiffusion | -4.77 | 53.84% | -3.51 | 27.45% | 4.5 |
| PepFlowww | -3.90 | 56.74% | -3.25 | 31.10% | 4.5 |
| PepGLAD | -6.11 | 60.27% | -5.14 | 39.83% | 3.0 |
| UniMoMo | -8.74 | 68.75% | -9.19 | 52.91% | 2.0 |
| SpecLig | **-12.91** | **75.43%** | **-16.39** | **75.00%** | **1.0** |

#### 8.2 Recovery related metrics

They measure the proportion of generated residues matching the reference peptide. **Amino Acid Recovery (AAR, ↑)** measures the proportion of generated residues matching the reference peptide. While multiple implementations exist, we adopt the most commonly used one in bioinformatics. As AAR is regarded unreliable, we only include it for completeness. **Complex RMSD (C-RMSD, ↓)** computes the RMSD of Cα atoms between generated and reference peptides after aligning by the target protein, while **Ligand RMSD (L-RMSD, ↓)** is similarly defined with alignment on the peptide itself.

Table S10. Complete comparison results for peptide design under Recovery category

| Method | AAR | C-RMSD | L-RMSD | **Rank** |
|---|---|---|---|---|
| RFDiffusion | 34.68% | 10.75 | 3.89 | 5.0 |
| PepFlowww | 35.47% | **7.57** | 3.57 | 3.0 |
| PepGLAD | **38.62%** | 8.13 | 2.86 | **2.0** |
| UniMoMo | 37.99% | 8.19 | 2.84 | 2.7 |
| SpecLig | 38.36% | 8.76 | **2.79** | 2.3 |

#### 8.3 Structural validity related metrics

**$Clash_{in}$ (↓)** and **$Clash_{out}$ (↓)** denote residue-level clashes within the peptide, and between the peptide and the target protein, respectively, defined by Cα atoms below 3.6574Å. **$JSD_{bb}$ (↓)** and **$JSD_{SC}$ (↓)** calculate Jensen-Shannon divergence between dihedral angle distributions of generated peptides and the dataset, for backbone (bb) and sidechain (sc) angles discretized into 10-degree bins.

Table S11. Complete comparison results for peptide design under Structural validity category

| Method | $Clash_{in}$ | $Clash_{out}$ | $JSD_{bb}$ | $JSD_{SC}$ | **Rank** |
|---|---|---|---|---|---|
| RFDiffusion | **0.06%** | 13.58% | 0.273 | 0.798 | 3.5 |
| PepFlowww | 2.72% | 4.62% | 0.240 | 0.693 | 3.8 |

| | | | | | |
|---|---|---|---|---|---|
| PepGLAD | 1.82% | 1.66% | 0.474 | 0.398 | 3.8 |
| UniMoMo | 0.57% | 1.55% | **0.237** | **0.182** | **1.8** |
| SpecLig | 0.33% | **1.26%** | 0.279 | 0.190 | 2.3 |

#### 8.4 Interaction related metrics

**ΔG(↓)** and **IMP(↑)** measure the binding energy calculated by pyRosetta, and the percentage of target proteins where generated peptides can outperform native binders. Besides, we denote **$\mathbf{JSD_{bs}}$ (↓)** as the Jensen–Shannon distance between the block distribution of the designed ligand and the block distribution of the reference ligand within the binding site.

Table S12. Complete comparison results for peptide design under Interaction category

| Method | ΔG | IMP | $JSD_{bs}$ | **Rank** |
|---|---|---|---|---|
| RFDiffusion | 92.215 | 5.38% | 0.212 | 5.0 |
| PepFlowww | 76.2776 | 14.13% | 0.129 | 4.0 |
| PepGLAD | 30.5195 | 17.2% | 0.014 | 2.6 |
| UniMoMo | 29.2101 | 40.71% | 0.022 | 2.3 |
| SpecLig | **-1.9218** | **43.94%** | **0.008** | **1.0** |

#### 8.5 Diversity related metrics

We calculate the ratio of unique clusters to total generations, with sequence identity above 40% and RMSD below 2Å as clustering thresholds.

Table S13. Complete comparison results for peptide design under Diversity category

| Method | Sequence | Structure | **Rank** |
|---|---|---|---|
| RFDiffusion | 0.155 | 0.616 | 4.5 |
| PepFlowww | 0.530 | 0.507 | 4.5 |
| PepGLAD | **0.687** | **0.698** | **1.0** |
| UniMoMo | 0.626 | 0.629 | 2.5 |
| SpecLig | 0.585 | 0.644 | 2.5 |

### 9 Hyperparameters

We provide the hyperparameters for SpecLig in Table S14.

Table S14. Hyperparameters of SpecLig in training and sampling

| Name | Value | Description |
|---|---|---|
| Variational AutoEncoder | | |
| epoch | 250 | Number of epochs to train. |
| warmup | 2000 | Number of warmup steps. |
| lr | 1e-4 | Learning rate. |
| embed size | 512 | Dimension of the element and block type embeddings. |
| hidden size | 512 | Dimension of hidden states. |
| edge size | 64 | Dimension of edge type embeddings. |
| k neighbors | 9 | Number of nearest neighbors in graph. |
| $L_{block}$ | 2 | Number of layers in the block-scale encoder and decoder. |
| $L_{struc}$ | 6 | Number of layers in the atom-scale encoder and decoder. |
| n_head | 8 | Number of heads for multi-head attention. |
| $\omega_1$ | 0.6 | The weight of KL divergence on the sequence. |
| $\omega_2$ | 0.8 | The weight of KL divergence on the structure. |
| $\omega_{\text{ac}}$ | 1.0 | The weight of atom coordinate MSE loss. |
| $\omega_{\text{bl}}$ | 1.0 | The weight of block type CE loss. |
| $\omega_{\text{tri}}$ | 1.0 | The weight of triplet loss. |
| $\omega_{\text{focal}}$ | 1.0 | The weight of focal atom-pair loss. |
| $\omega_{\text{cen}}$ | 0.2 | The weight of block-centroid position loss. |
| $\omega_{\text{bond}}$ | 0.5 | The weight of bond prediction loss. |
| $\omega_{\text{dist}}$ | 0.5 | The weight of local distance loss. |
| Latent Diffusion Model | | |
| T | 100 | Number of total steps for diffusion. |
| hidden size | 512 | Dimension of hidden states. |
| n_layers | 6 | Number of layers in noise-prediction network. |
| n_head | 8 | Number of heads for multi-head attention. |
| $\omega_{\text{lpl}}$ | 2.0 | The weight of LPL loss. |
| ω | 1.0 | The weight of noise prediction. |
| $\tau_{temp}$ | 1.2 | Temperature-scaled normalization factor. |
| $G_{\text{set}}$ | 4.0 | Clipped bound for norm of gradient. |
| guidance_scale | 3.4 for peptides / 1.8 for small molecules | The weight of energy guidance for different tasks. |

### 10 Ablation Experiments

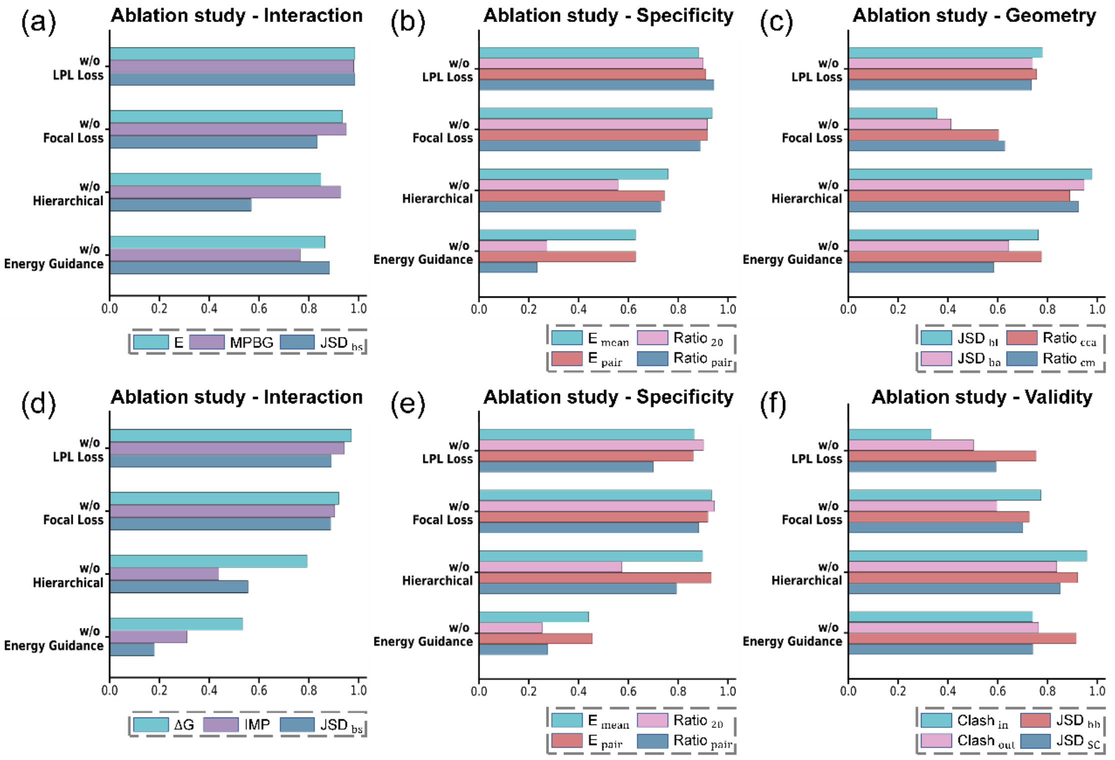


Figure S1. Ablation studies of SpecLig components. In small molecule design, we systematically evaluate the impact of latent perceptual loss (LPL), focal loss, hierarchical network architecture, and energy-guided sampling on interaction (a), target specificity (b), and geometric validity (c) categories. In peptide design, we evaluate the same modules on interaction (d), target specificity (e), and structural validity (f) categories. Given that the value ranges of different evaluation metrics vary significantly, we utilized relative proportions for clarity.

We conducted comprehensive ablation studies on SpecLig's architectural components, evaluating their performance separately on small molecule design and short peptide design tasks. Our analysis particularly focuses on quantifying each module's contribution to binding affinity, molecular specificity, and conformational validity. As evidenced in Figure S1, energy-guided sampling emerges as the critical factor that simultaneously enhances both binding affinity and target specificity across both molecular design domains. The designed loss functions effectively bridge the representational gap between the variational autoencoder (VAE) and latent diffusion model (LDM), thereby preserving structural consistency in the generated molecules, as reflected in their significant contributions to geometric validity metrics.

#### 11 Self-Adaptive Molecular Sizes

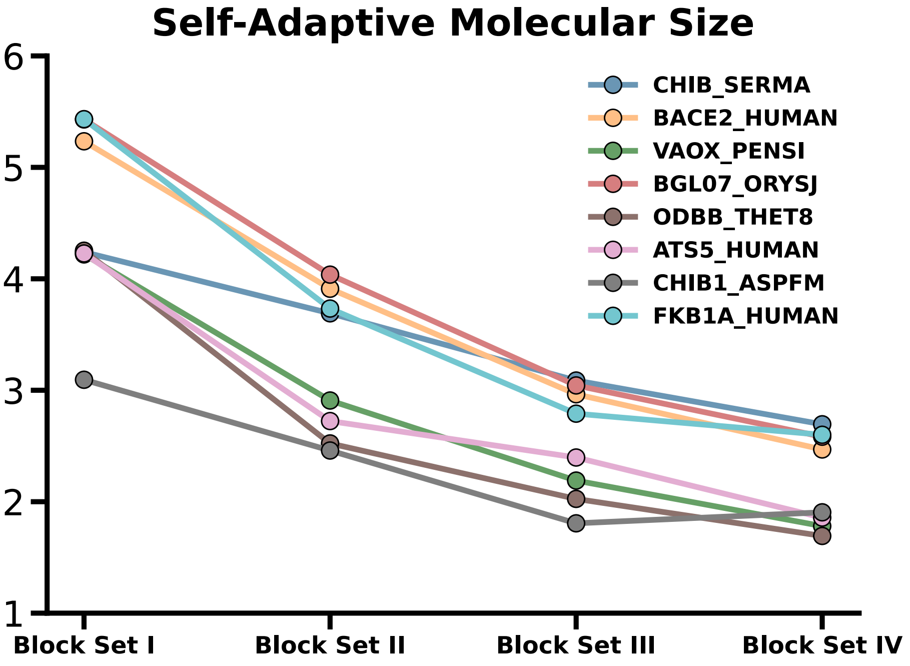


Figure S2. Comparisons of number of atoms in each block for small molecules under different settings of designated number of blocks.

Inspired by observations of context-aware generation behavior in SBDD, we investigated whether SpecLig exhibits similar adaptive capabilities when designing small molecules under varying structural constraints. For eight representative protein targets (shown in Figure S2), we generated 2,000 ligands per target with randomly assigned block counts ranging from 3 to 22. When analyzing the average atoms per block across four block-count categories (3–7, 8–12, 13–17, and 18–22 blocks), we observed a clear inverse relationship between block quantity and average atoms. With fewer blocks, SpecLig automatically selected larger molecular fragments to maintain sufficient binding interface coverage. As block count increased, the model progressively favored smaller fragments to prevent steric clashes. This self-adaptive behavior demonstrates SpecLig's capacity to dynamically adjust its generation strategy based on structural constraints.

### 12 Case Study

In Figures S3 and S4, we present SpecLig's performance on eight additional targets, demonstrating how our method enhances both binding affinity and specificity relative to native ligands, separately for small-molecule and short-peptide design.

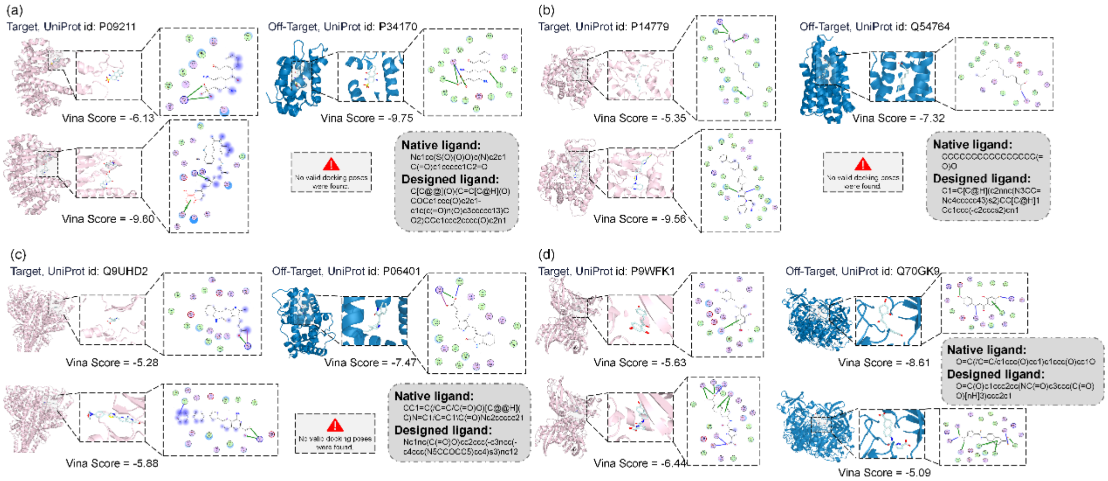


Figure S3. For each subplot, we present a pairwise docking analysis comparing a target protein (pink) and a non-target protein (blue). The top row shows the binding pose and interactions of the native ligand, while the bottom row displays those of the ligand designed by SpecLig. Interaction profiles were generated using Molecular Operating Environment (MOE). The corresponding SMILES strings are provided in gray boxes for reference. The target and non-target pairs used in small molecule design are: Glutathione S-transferase P and GOBP2 (a), Cytochrome P450BM-3 and Aldehyde decarbonylase (b), threonine-protein kinase TBK1 and Progesterone receptor (c), and UPF0336 protein Rv0635 and Fluorinase (d).

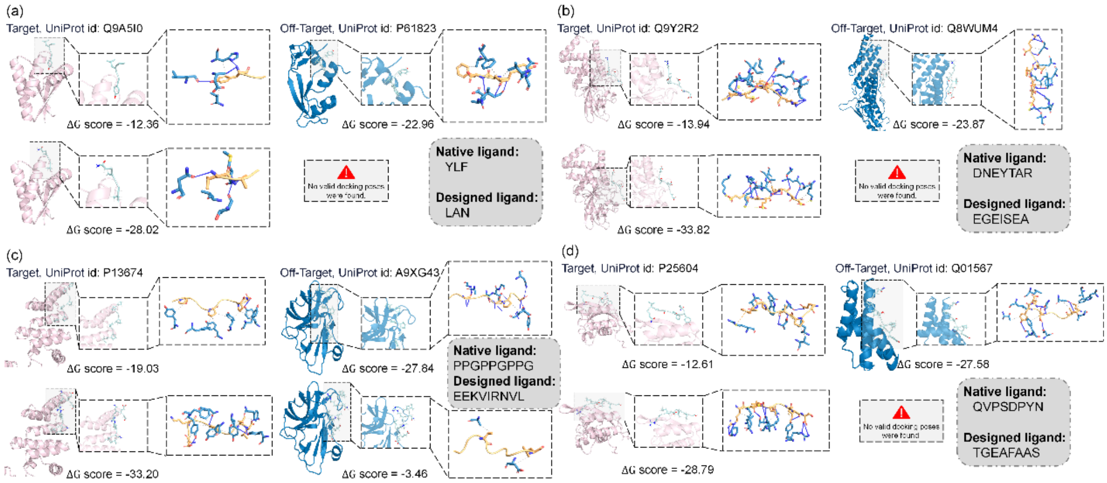


Figure S4. For each subplot, we present a pairwise docking analysis comparing a target protein (pink) and a non-target protein (blue). The top row shows the binding pose and interactions of the native ligand, while the bottom row displays those of the ligand designed by SpecLig. Interaction profiles were generated using Rosetta. The corresponding ligand sequences are provided in gray boxes for reference. The target and non-target pairs used in peptide design are: ATP-dependent Clp protease adapter protein ClpS and Ribonuclease pancreatic (a), Tyrosine-protein phosphatase non-receptor type 22 and PDCD6-interacting protein (b), 4-PH alpha-1 and Genome polyprotein (c), and ESCRT-I complex subunit VPS23 and Pilotin OutS (d).

#### 13 Sensitivity Analysis of Energy Guidance

We systematically investigate the impact of energy guidance strength on binding affinity, target specificity, and sequence diversity in peptide design. As demonstrated in Figure S5, a pronounced diversity-specificity and diversity-affinity trade-off emerges: increasing energy guidance progressively reduces sequence diversity, while affinity and specificity initially improve but decline under excessive guidance. Strong energy guidance enforces convergence toward fixed motifs, substantially constraining conformational exploration. We further attribute the late-stage degradation in affinity and specificity to near-mode-collapse behavior in generative modeling—overly constrained gradients concentrate probability mass on sparse local optima, thereby suppressing beneficial exploration of high-potential sequence space.

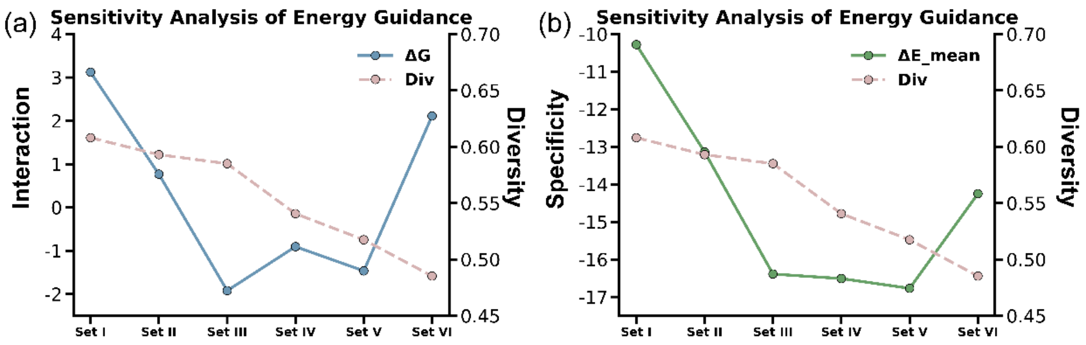


Figure S5. Design performance on the test set across energy guidance strengths (predefined weights: [2.6, 3.0, 3.4, 3.8, 4.2, 4.6]). (a) Trade-off between binding affinity (ΔG) and sequence diversity; (b) Variation in target specificity ($\Delta E_{mean}$) versus sequence diversity.
